# A CELL BIOLOGY STUDY REVEALS NEW INSIGHTS INTO THE TRANSPORT MECHANISMS OF OOMYCETE EFFECTORS

**DOI:** 10.1101/2024.10.29.620966

**Authors:** Iga Tomczynska, Michael Stumpe, Didier Reinhardt, Markus M. Geisler

## Abstract

Oomycete pathogens use dedicated effector proteins (also known as virulence factors) that manipulate the host during infection to the benefit of the pathogen. Despite extensive efforts, the understanding of how oomycete effector proteins are transported and delivered to the host is limited. Here, we show that *Phytophthora capsici* infecting *Nicotiana benthamiana* secretes its effectors through secretory pockets, haustoria, and from hyphae. We provide evidence that Avr3, belonging to the class of RxLR effector proteins that can be secreted and translocated to the plant cell, is found to accumulate in a distinct ring-shaped zone around the neck region of haustoria. The association with the haustorial neck is determined by an RxLR-EER motif and mutating this motif redirects Avr3 to the extrahaustorial matrix enveloping the full haustoria body. Furthermore, mass spectrometry analysis of immunoprecipitated Avr3 indicates that the RXLR motif remains intact when Avr3 is secreted from the haustorium during plant infection. We thus propose that the RXLR-EER motif determines the entry in the host cell by guiding Avr3 to the haustorial neck region, which may be a structure analogous to the Biotrophic Interfacial Complex of the fungus *Magnaporthe* sp.

## INTRODUCTION

The oomycete genus Phytophthora is represented by over 200 species. The vast majority of them, like *P. infestans* or *P. capsici*, are plant pathogens and due to their ecological, economic, social, and scientific impact, they remain in the spotlight of plant pathology (Brasier *et al*., 2022). In order to overcome plant defense and to achieve infection, Phytophthora produces and secretes a suite of proteins called effectors (or virulence factors) that facilitate colonization of the host plant. Considering their site of action, two types of effectors can be distinguished. The first group comprises proteins that are secreted and carry out their functions in the apoplastic space. They include hydrolytic enzymes degrading plant cell walls, such as xyloglucanase XEG1 (Ma *et al*., 2015), elicitins that are oomycete specific extracellular sterol carriers (Vauthrin *et al*., 1999; Wang, W *et al*., 2021) or Nep1-like proteins (NLPs) that are best known for their cytotoxic effect on the plant membrane (Oome & Van den Ackerveken, 2014). Apart from these intensively studied examples, a large portion of the Phytophthora proteome has been predicted to be extracellular based on the presence of a signal peptide sequence but direct functional characterization is lacking in most predicted effectors (Raffaele *et al*., 2010). One such example is Phytophthora conserved protein PITG_02399 (in *P. infestans*)/ Pc508140 (in *P. capsici*).

The second group includes intracellular RxLR effectors. Upon translocation to the plant cell, they bind to host proteins (targets) and impair their function to affect plant defense. The name of this group stems from the characteristic amino acid motif RxLR (Arg, any, Leu, Arg) that is located downstream of the signal peptide and often followed by an E/D-rich domain (Glu/Asp). This motif was shown to be implicated in effector delivery to the plant cell (Whisson *et al*., 2007; Dou *et al*., 2008). While the RxLR motif is conserved and can be used as a hallmark for protein annotation, the overall sequence homology between RxLR effectors is rather low (Jiang *et al*., 2008). One rare example of a conserved RxLR effector is *P. capsici* RxLR7 (Chen *et al*., 2014). Additionally, one of the most extensively studied RxLR effectors, Avr3a of *P. infestans*, has homologs in *P. sojae* and *P. capsici* (Bos *et al*., 2006; Li *et al*., 2019).

The main site of protein secretion during infection by Phytophthora is the haustorium (Wang *et al*., 2017; Wang *et al*., 2018; Kagda *et al*., 2020), a structure that enters a living plant cell and forms a pathogen-host interface. Lying outside haustorial cell wall is the extrahaustorial matrix (EHMx; equivalent of modified apoplast), which is surrounded by an extrahaustorial membrane (EHM; modified plant plasma membrane) (Enkerli *et al*., 1997b). Haustorial expansion also triggers one of the earliest plant defense responses against penetrating pathogen. Deposition of callose papillae at the haustorium serves as a mechanical barrier against the invader (Wang, Y *et al*., 2021).

So far, most of the scientific effort in the Phytophthora field has focused on RxLR effector-host target interaction and its consequences for plant immunity. Studies relied on heterologous expression of effectors in planta and aim to mimic conditions when the effector is already present in the host tissue (Tomczynska *et al*., 2018; Tomczynska *et al*., 2020). There is however a substantial gap in the knowledge on how Phytophthora virulence factors are transported. Only a few studies have investigated the secretion of apoplastic effectors by the pathogen (Wang *et al*., 2017; Wang *et al*., 2018; Kagda *et al*., 2020). Much more attention has focused on RxLR effector delivery to the plant symplast, although this topic is also surrounded by controversy related to conflicting data.

The RxLR-EER motif of *P. infestans* Avr3a was shown to be essential for effector translocation into the plant cell (Whisson *et al*., 2007). Following this discovery, two contradictory models were proposed to explain this phenomenon. According to the first one, effector entry to the host cell is pathogen-independent and occurs via binding of the RxLR motif to phosphoinositol-3-phosphate (PI3P) exposed on the surface of eukaryotic cells (Dou *et al*., 2008; Kale *et al*., 2010). Later, this model was criticized for issues with the experimental system (Wawra *et al*., 2013) and reports showed that, in absence of pathogen, host cells alone do not have a mechanism that can mediate effector uptake (Petre *et al*., 2016; Wang *et al*., 2017). The second model proposed that the RxLR motif is cleaved off prior to secretion, therefore serving as a sorting signal for a specific route of effector transport inside the hyphae (Wawra *et al*., 2017).

The major limitation of prior studies (Dou *et al*., 2008; Kale *et al*., 2010; Wawra *et al*., 2017) lies in their experimental systems, which rely exclusively on either pathogen or plant cells, without examining the interaction between pathogen and plant cells simultaneously. Given that the haustorium develops exclusively during the biotrophic phase of infection and serves as the major secretory site for virulence factors (Wang *et al*., 2017; Wang *et al*., 2018; Kagda *et al*., 2020), we aimed to investigate previous uncertainties by studying Phytophthora effector transport in the presence of haustoria. Previous studies have shown that in contrast to apoplastic effectors, RxLR effectors are transported to haustoria by a Brefeldin A-resistant pathway (Wang *et al*., 2017; Wang *et al*., 2018) and associate with plant proteins that promote clathrin-mediated endocytosis (Wang *et al*., 2023). As far as we know, despite discussions on the role of the ER and secretory vesicles (Boevink *et al*., 2020), there is not yet conclusive evidence for the transport of secreted proteins from the hyphae lumen to haustoria. Whisson et al. (2007) showed that both wild-type Avr3a and a mutated version lacking the RxLR-EER motif reach haustoria, but only the wild-type is translocated to the host cell. This raises the question of what differentiates their transfer from haustoria to plant cells if both proteins reach the secretory site. Additionally, it remains unclear how the secretory site is established over time.

To address the issues outlined above, we examined the transport of seven proteins previously predicted or identified as effectors secreted by Phytophthora. These were *P. capsici* RxLR effectors Avr3 and RxLR7, *P. sojae* Avh110 and apoplastic proteins elicitin, NLP, XEG1 and Pc508140. Our cell-biological study performed on a wide range of *P. capsici* transformants revealed that transport of both apoplastic and RxLR effectors in the hyphae relies on mobile structures that can be secretory vesicles. We observed that haustorium serves as a main secretory site, however we noted two exceptions: XEG1 was secreted also from hyphal regions whereas a cell-death inducing effector RxLR7 was absent in haustoria. Additionally, we observed secretory pockets functioning as initial secretion sites before haustoria maturation. Consistent with findings in *Magnaporthe oryzae* (Khang *et al*., 2010), our results show distinct accumulation regions for two classes of *P. capsici* effectors at secretory sites. Apoplastic proteins are delivered into EHMx, whereas the RxLR effector Avr3 accumulates at a distinct region within the cell wall at the neck of *P. capsici* haustoria. We also provide strong evidence that Avr3 localization at haustorial neck region depends on the RxLR-EER motif fragment, supported by MS analysis showing that the RxLR motif of Avr3 remains intact during infection. Taken together, our discoveries provide a significant contribution to understand the complexity of Phytophthora effector transport.

## RESULTS

### Cell wall staining as a strategy to reveal the organization of *P. capsici* haustoria

Previous fluorescence microscopy studies used free tags to label the Phytophthora cytoplasm for co-localization experiments but they did not provide an ideal reference for investigating the organization of secretory sites (Wang *et al*., 2017; Wang *et al*., 2018; Kagda *et al*., 2020). Both the EHMx and the plasma membrane of the haustorium are located “outside” the cytosol, therefore the free fluorescent tag does not help to distinguish their localization. In this work, we employed the Phytophthora cell wall as a marker for subsequent co-localization studies. In oomycetes, β-1,3-glucans are detectable at all stages of haustorial development (Enkerli *et al*., 1997b; Mims *et al*., 2004). Since structural components of the *P. infestans* cell wall, callose and cellulose (Hachler & Hohl, 1982), can be stained by aniline blue and calcofluor white, respectively, we aimed to use dyes to visualize hyphae and haustoria. Both dyes vary in emission spectra and specificity. Calcofluor white can be used with GFP, YFP, mVenus, RFP and mCherry (Ursache *et al*., 2018) but aniline blue, due to the fact that its fluorescence can be shifted up to a wavelength of 506 nm (Smith & McCully, 1978), has a narrower range of compatibility with fluorescent tags. Cellulose is also a plant cell wall component (Ursache *et al*., 2018) and callose can be deposited by the plant at infection sites as a response against pathogen penetration (Enkerli *et al*., 1997a; Wang, Y *et al*., 2021). Therefore, the potential drawback of this approach is that the signal derived from Phytophthora cell walls could be masked by signal from the plant cell wall.

First, the utility of aniline blue and calcofluor white as tools to label Phytopthora cell wall was evaluated on *P. capsici* transformants expressing a secreted mCherry construct (Supplemental Fig. S1 and S2A). Interestingly, without penetration of the host tissue, mCherry fused with a signal peptide sequence expressed by *P. capsici* was retained inside the hyphae (Supplemental Fig. S1A). Hence, in the absence of an evident secretory site, non-penetrating hyphae were excluded for further observation and our attention was restricted to penetrating hyphae during infection, given their notable accumulation of secreted mCherry around haustoria (Supplemental Fig. S1B).

Then, the stained Phytophthora cell wall as a marker for co-localization experiments was further validated with *P. capsici* transformants that expressed mCherry constructs localized to the following subcellular compartments: the extrahaustorial matrix (EHMx, mCherry fused with signal peptide) (Supplemental Fig. S2A), the haustorial symplast (endoplasmic reticulum, mCherry fused with KDEL motif) (Supplemental Fig. S2B), and the haustorial membrane (mCherry fused with myristoylation signal) (Supplemental Fig. S2C). The correct orientation between the examined sites and the haustorial cell wall was confirmed through signal intensity profiles (Supplemental Fig. S2D).

Interestingly, there was a clear difference in the appearance of signal derived from the haustorial plasma membrane and EHMx. While the plasma membrane displayed a shape of a continuous line (Supplemental Fig. S2C and E), EHMx filled with SP-mCherry appear as a shape with protuberances (Supplemental Fig. S1B and S2A) which is consistent with electron microscopy pictures of convoluted EHM (Enkerli *et al*., 1997b).

The sites where the pathogen penetrated the plant cell wall and entered the cells through cell wall perforations were clearly visible in the tissue infected with *P. capsici* transformants expressing the plasma membrane marker. Interestingly, for some of perforation points, we could not pinpoint corresponding haustoria responsible for their creation. Instead, we observed perforations that seemed to be located on the hyphae-plant cell wall surface (Supplemental Fig. S2E and S3). This led us to conclude that also in the absence of a haustorium, Phytophthora can release lytic enzymes to digest the host cell wall.

### Phytophthora effectors are transported to secretory pockets

To further address the phenomenon of plant cell wall perforations in the absence of haustoria, we generated a set of *P. capsici* transformants expressing tagged versions of proteins secreted to the apoplast (elicitin, NLP, Pc508140) and RxLR effectors (Avr3, Avh110). In *P. capsici* transformants infecting plant tissue, both types of effectors accumulated not only in haustoria but also in flat structures adjacent to the hyphae, which we further refer to as “secretory pockets” (Fig. 1). Unlike haustoria, which are characterized by their finger-like protrusions, secretory pockets lack a distinct dimensional structure, since they can hardly be distinguished in brightfield images (Fig. 1A and B). Only at high magnification, secretory pockets can be identified as the space positioned adjacent to the hyphal surface, containing deposited secretory material (Fig. 1C).

**Figure 1.**
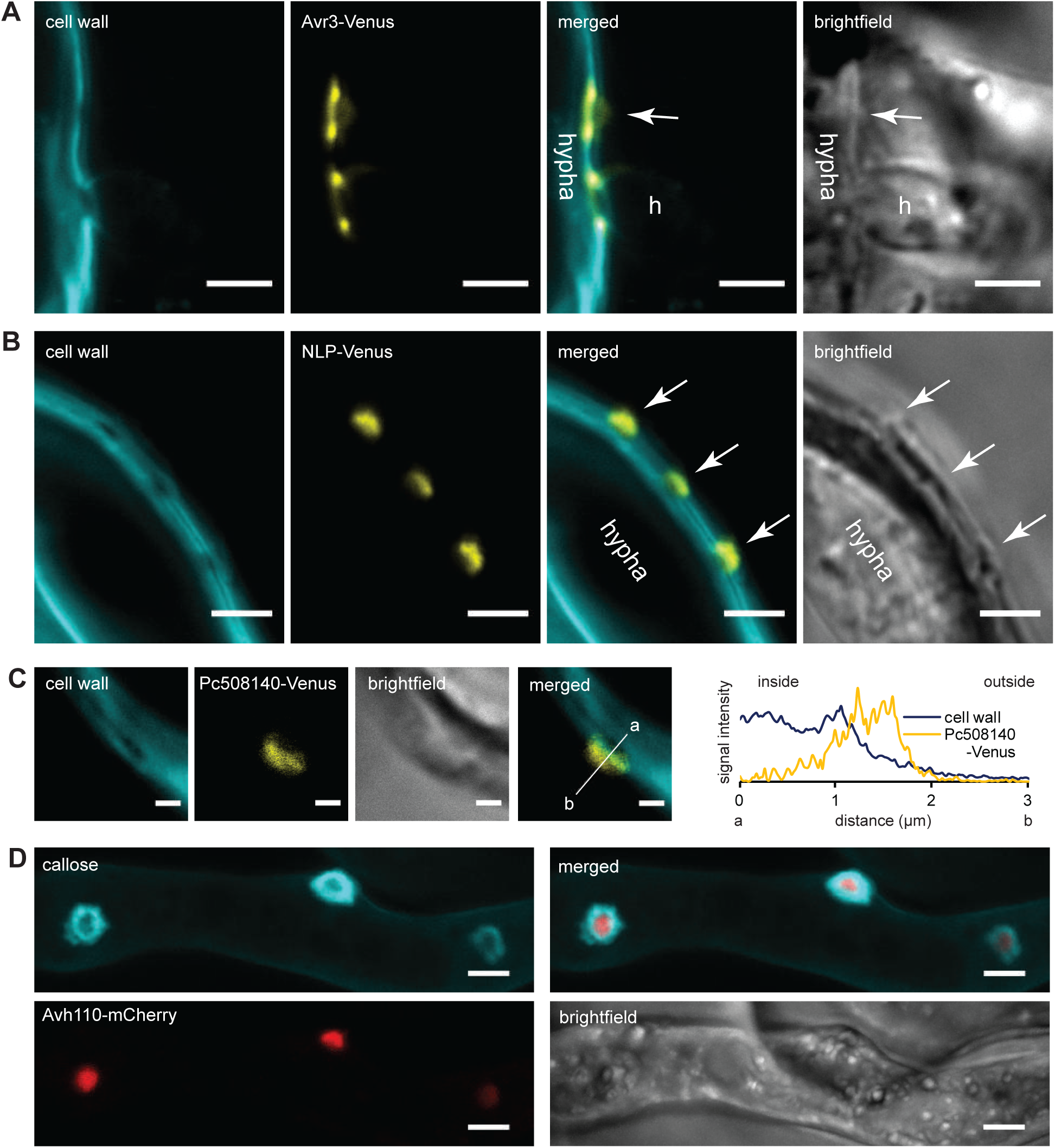
*P. capsici* secretes effectors to secretory pockets. Representative confocal microscopy pictures of cell wall stained with calcofluor white and (**A**) the secretory sites for RxLR effector Avr3-Venus: the secretory pocket marked with an arrow vs. the mature haustorium (h), scale bar 3μm. (**B**) the secretory pocket filled with NLP-Venus along the hypha, position of the secretory pocket on the hyphal surface marked with arrows, scale bar 3μm. (**C**) the secretory pocket filled with Pc508140-Venus, scale bar 1μm. The picture is accompanied by the relative fluorescence intensity plot of the signals measured along the line shown in merged picture (a-inside, b-outside hypha). (**D**) Representative confocal microscopy pictures (maximal projection z-stack of 10 slices) of callose deposits stained with aniline blue around Avh110-mCherry positive secretory pockets, scale bar 3μm.

In numerous diseases caused by oomycetes, transmission electron micrographs showed the presence of a matrix that initially appears as an electron-dense deposit of secreted material covered by the host plasma membrane and separated from the pathogen hypha by the host cell wall (Chou, 1970; Enkerli *et al*., 1997b). Our observations of frequent accumulation of apoplastic and intracellular RxLR effectors in the secretory pockets are reminiscent of these descriptions (Supplemental Fig. S4). Moreover, similarly to the reaction against haustoria, we noted that plant cells formed callose-containing papillae in reaction to the secretory pockets indicating that the host can detect and respond to this type of secretory site (Fig. 1D). According to the literature, deposits of secretory material represent sites from which new haustoria emerge and subsequently develop to the mature stage. After the pathogen breaks through the host cell wall, the matrix filled with secretory material likely extends across the surface of the developing haustorium that grows into the plant cell lumen and invaginates the host-cell plasma membrane (Enkerli *et al*., 1997b).

While technical constraints prevented us from conducting a time-lapse series showcasing the development of individual secretory pockets, the range of intermediate stages between the secretory pocket and the haustorium in our study (Supplemental Fig. S5) aligns with the conclusion made by previous authors (Chou, 1970; Enkerli *et al*., 1997b). Our observation of secretory sites indicates that *P. capsici* secretes proteins prior to haustorium formation, and that secretory pockets serve to prepare future infection sites for invasion and haustorium development. These secretory sites remain functional, at least for elicitin secretion, in fully expanded haustoria (Supplemental Fig. S5E).

### Phytophthora apoplastic proteins reach their secretory sites via vesicle-mediated transport

It has been commonly accepted that haustoria serve as sites of active secretion, potentially for any protein with a signal peptide (Kagda *et al*., 2020). Therefore, following our discovery of secretory pockets, we subsequently investigated the presence of other target sites to which apoplastic proteins (elicitin, Pc508140, NLP, and XEG1) are directed.

All the examined proteins were found to accumulate at haustoria (Supplemental Fig. S6). Intensity profiles of the fluorescent signals derived from elicitin, Pc508140 and NLP pointed out that they are localized outside the haustorial cell wall which proves their secretion to the EHMx (Fig. 2A-C). Notably, we found haustoria where NLP-Venus was not detected (Supplemental Fig. S6C). Inside the hyphae, we observed the fluorescent signal of secreted proteins within spherical, mobile structures of approximately 1 μm diameter (Supplemental Movie S1 and S2) which suggests that transport towards haustoria is mediated by secretory vesicles. This finding aligns with a previous study that reported transport of apoplastic virulence factors proceeding via the conventional ER-Golgi dependent secretory pathway (Wang *et al*., 2017). We also confirm secretion of the conserved Phytophthora effector Pc508140. In *P. infestans*, the homologue PITG_02399 is up-regulated during infection of tomato (Zuluaga *et al*., 2016) and predicted to be secreted (Raffaele *et al*., 2010) but was not detected among extracellular proteins when mycelium was grown in liquid media (Meijer *et al*., 2014), suggesting that this protein is only secreted upon contact with host tissues.

**Figure 2.**
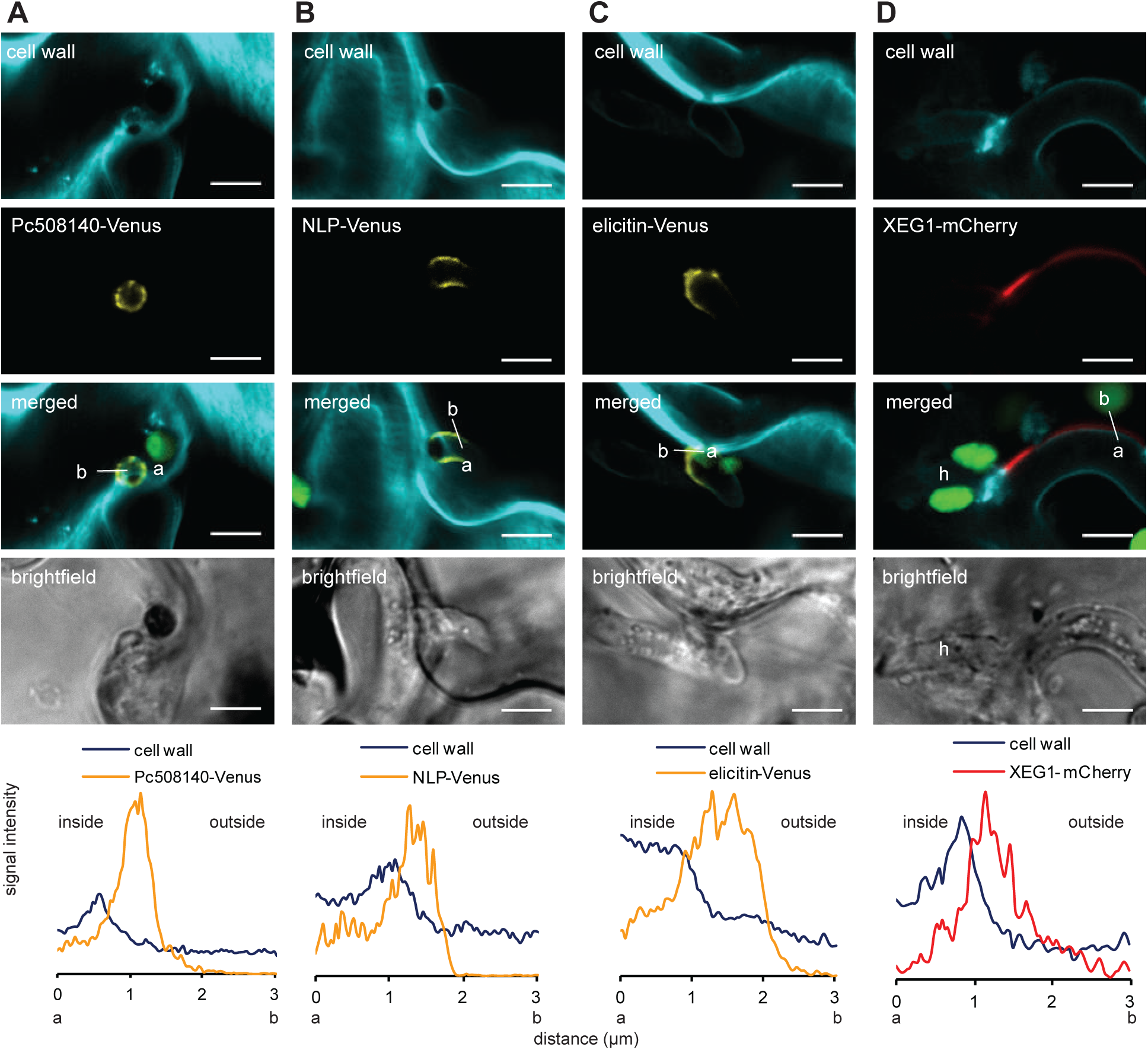
*P. capsici* effectors reach the extrahaustorial matrix and apoplast. Representative confocal microscopy pictures of (**A**) Pc508140-Venus, (**B)** NLP-Venus, (**C**) elicitin-Venus secreted outside haustorium. Cell wall stained with calcofluor white, scale bar 5 μm. (**D**) Confocal microscopy pictures (maximal projection z-stack of 10 slices) of XEG1-mCherry secreted outside hyphae. Haustorium structure (h) distinguished by the callose collar stained with aniline blue, scale bar 5 μm. Chloroplasts labeled in green. The bottom panels show relative fluorescence intensity plots of signals measured along the line shown in merged picture (a-inside, b-outside haustorium) on respective merged pictures.

Interestingly, we observed that haustoria are not the sole secretory site. XEG1 turned out to be an exception, accumulating not only at haustoria (Supplemental Fig. S6) but also in the extracellular space around hyphae (Fig. 2D), potentially reflecting its function. Conceivably, the xyloglucanase encoded by *XEG1* (Ma *et al*., 2015) allows the pathogen to pass in the apoplast between host cells by degrading or softening the host hemicelluloses while exploring the apoplastic space. Indeed, previous research demonstrated that filamentous fungi can secrete digestive enzymes to their environment from the hyphae (Read, 2011). The absence of XEG1 signal from some haustoria, and its presence outside the hyphal surface (Fig. 2D) indicates that XEG1 is transported by a different secretory pathway than other apoplastic effectors such as elicitin, Pc508140, and NLP. Alternatively, the explanation for the “empty” haustoria we observed, including in the NLP-Venus transformants (Supplemental Fig. S6C), could be that they are dysfunctional, with their secretory function aborted.

### RxLR effector Avr3 is translocated to the plant cell whereas RxLR7 is retained in Phytophthora nuclei

To further explore the secretory transport in Phytophthora during infection, we focused on intracellular RxLR effectors. Tagged RxLR effectors Avr3, RxLR7 and Avh110 were transformed into *P. capsici*. Whereas Avr3 and RxLR7 originate from *P. capsici*, Avh110 is a *P. sojae* effector and shares low amino acid sequence identity (below 25%) with the closest homologue from *P. capsici* (Qiu *et al*., 2023). Inside the hyphae growing in the plant tissue, the signals derived from Avr3 and Avh110 are present in the dotted structures that could be transport vesicles and accumulate at the haustoria, which indicates their successful delivery to the secretory site (Fig. 3A and B, C). Notably, Avr3 and Avh110 accumulated mainly at the haustorial base and not at the protuberances of EHMx. By contrast, RxLR7 labeled irregularly shaped structures inside the hyphae, and we did not observe any secretory sites for RxLR7 (Fig. 3C). According to the prediction with NLSdb (Nair *et al*., 2003), RxLR7 carries a nuclear localization signal (NLS) between amino acids 229 and 233 (’RTRQR’). We thus hypothesized that RxLR7 instead of being secreted, is recruited to *P. capsici* nuclei. Co-localization between RxLR7-Venus and *P. capsici* nucleolar marker fibrillarin (FiB-mCherry) (Fang *et al*., 2017) shows that RxLR7 indeed localizes to the nuclei of *P. capsici* (Fig. 4A).

**Figure 3.**
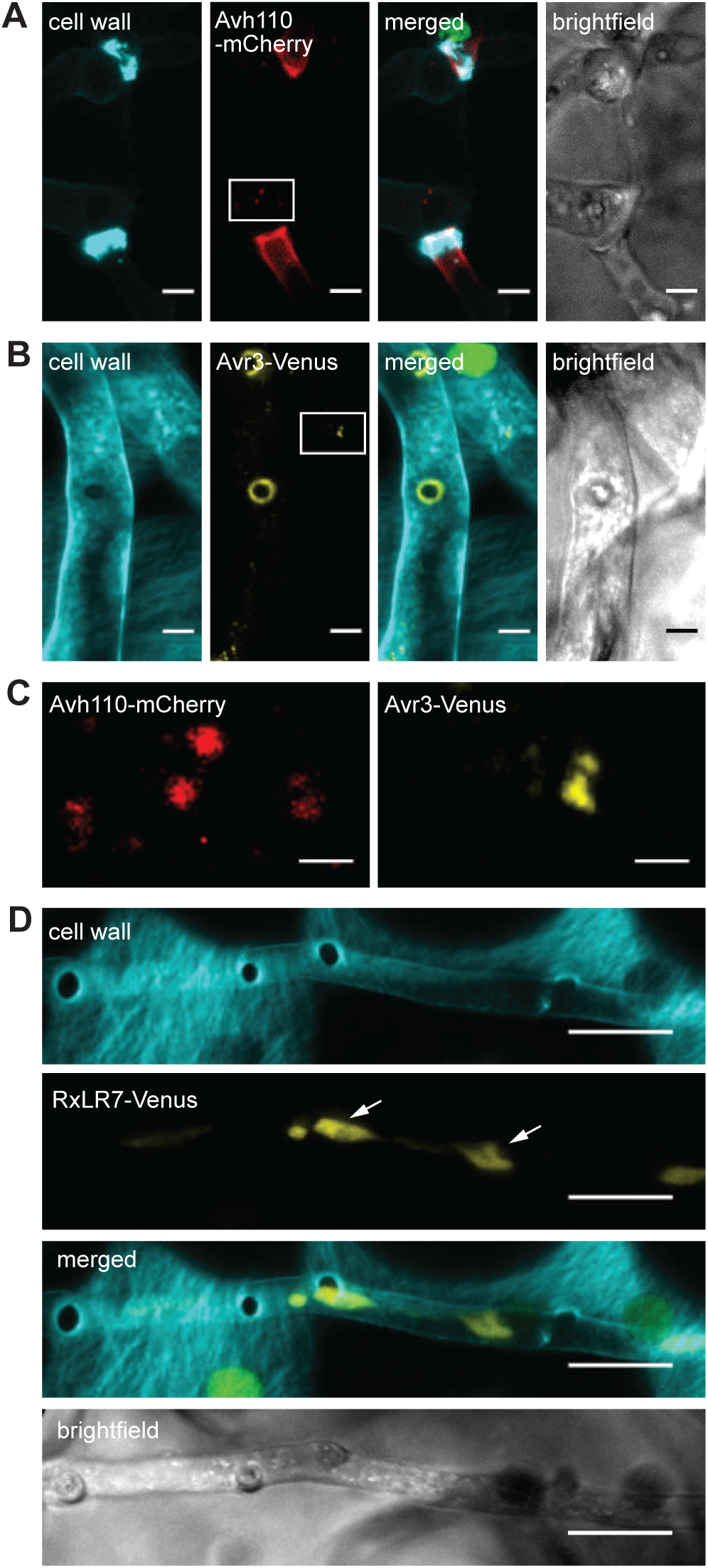
RxLR effectors Avr3 and Avh110 but not RxLR7 are delivered to haustoria via vesicle-like structures. Representative confocal microscopy pictures of *P. capsici* expressing (**A**) Avh110-mCherry (maximal projection z-stack of 12 slices), haustoria can be distinguished by the presence of callose collar, scale bars 3 μm. (**B**) Avr3-Venus (maximal projection z-stack of 11 slices), scale bars 3 μm. (**C**) Close-up view of the white boxes marked in **A** and **B** show dotted structures that could be transport vesicles in the hyphae, scale bars 1 μm. (**D**) RxLR7-Venus (maximal projection z-stack of 11 slices), scale bar 10 μm. The arrow marks RxLR7-positive organelles inside the hypha. Cell wall stained with aniline blue (**A**) or calcofluor white (**B** and **D**). Chloroplasts labeled in green.

**Figure 4.**
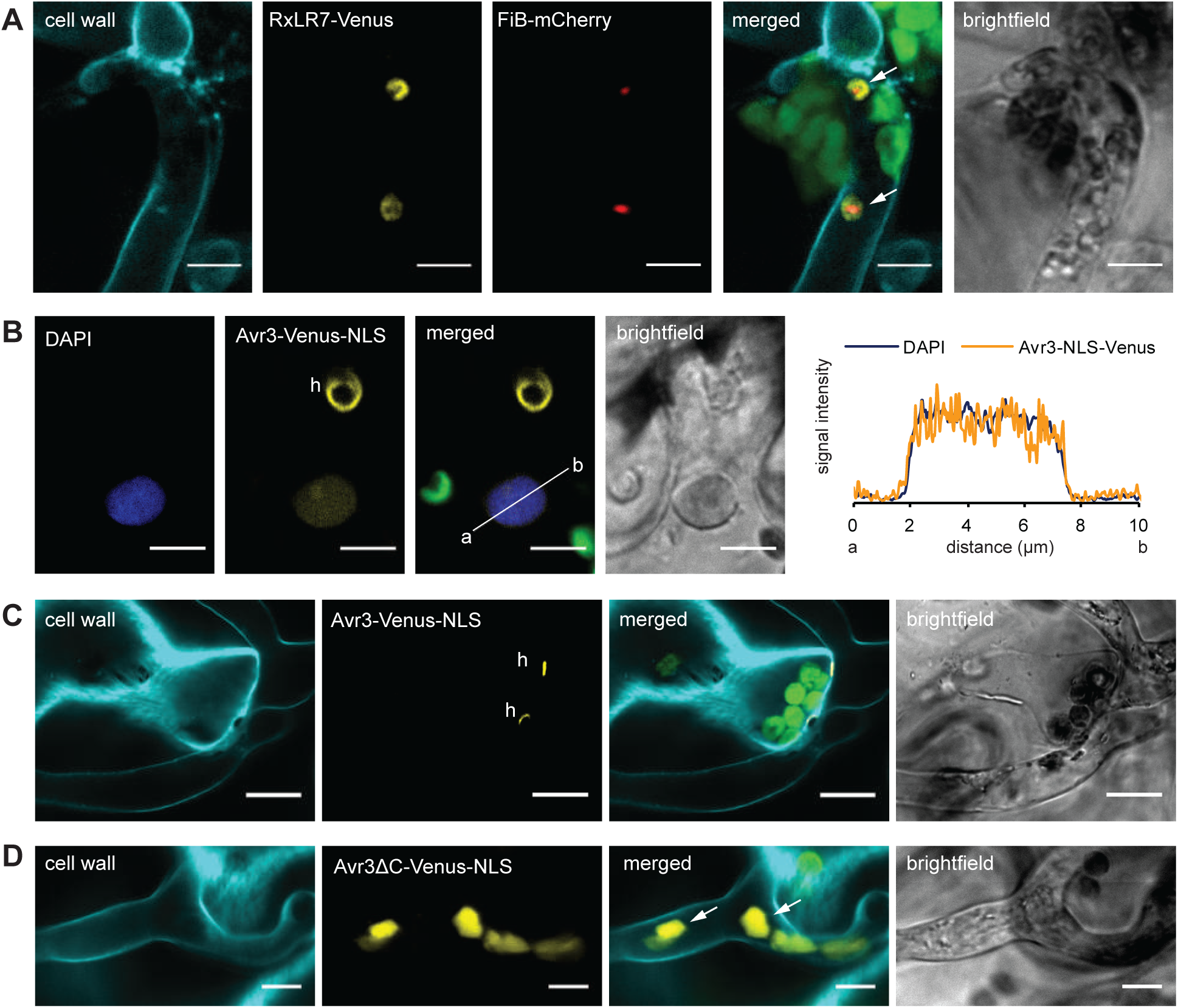
RxLR7 is retained in *P. capsici* nuclei whereas Avr3 is translocated to the host cell. Representative confocal microscopy pictures of (**A**) RxLR7-Venus co-localizing with the nucleolar marker FiB-mCherry inside *P. capsici* hyphae, scale bar 5 μm, (**B**) NLS fusion of Avr3-Venus present in haustorium (h) and in plant nucleus of the host cell (stained with DAPI) penetrated by this haustorium. The picture accompanied by relative fluorescence intensity plot of both Venus and DAPI signals detected along the line from the point a to b shown in the merged picture, scale bar 5 μm. Confocal microscopy pictures showing localization of full length Avr3 (**C**) and its truncated version (**D**) fused with Venus-NLS. Avr3-Venus-NLS construct expressed by *P. capsici* is transported to haustoria (h) whereas its truncated version that does not carry the sequence downstream of EER motif is targeted to the nuclei (marked with arrows in the merged picture). Scale bars 10 μm (**C**) 5 μm (**D**) respectively. Cell wall stained with aniline blue (**A**) or calcofluor white (**C** and **D**). Chloroplasts labeled in green.

In our study, RxLR effectors Avr3 and Avh110 localized to a population of the dotted structures within the hyphae. Although their role remains unknown, prior literature suggests that they could be involved in an unconventional secretory pathway (Wang *et al*., 2017; Wang *et al*., 2018). In a previous study, RxLR7 was identified as an RXLR effector with highly conserved orthologues (>70% amino acid identity) across oomycetes and the allele from *P. capsici* triggered cell death when ectopically expressed in *N. benthamiana* (Chen *et al*., 2014). Strikingly, in our experiments, RxLR7 was not secreted by *P. capsici* transformants but directed to the nuclei inside the pathogen’s hyphae.

The prediction of RxLR effectors relies on the RxLR motif accompanied by a signal peptide that determines secretion. While numerous studies disqualify a protein candidate as a secreted RxLR effector if it carries also a KDEL/HDEL motif responsible for ER retention (Evangelisti *et al*., 2017; Rojas-Estevez *et al*., 2020), the presence of an NLS is considered as a characteristic feature for effectors targeting the plant nucleus rather than as a factor restricting their secretion (Rivas & Genin, 2011). On the other hand, literature provides examples that a signal peptide is not dominant over an NLS motif (Kiefer *et al*., 1994) and RxLR7 might be one of them.

As discussed by other authors (Wawra *et al*., 2013), it is not clear how to distinguish functionally relevant RxLR effectors from proteins that contain an RxLR motif by coincidence. Although we cannot exclude that RxLR7 secretion may occur under certain infection conditions (not provided in our experiments), or by different *P.capsici* isolates, the fact that RxLR7 was found to be retained in the hyphae questions whether this protein can be considered a genuine RxLR effector. It also shows that classifying proteins as effectors based solely on bioinformatic prediction and amino acid motifs requires additional functional evaluation.

Artificial fusion of an NLS to the effector has been demonstrated as a technique to concentrate a pathogen protein within the plant nucleus, commonly used to verify effector translocation to the plant cell (Khang *et al*., 2010). The fusion of Avr3 with SV40 NLS (Simian virus 40 T antigen nuclear localization sequence) confirmed that Avr3 is indeed transported to the plant cell, as indicated by the colocalization of Avr3-Venus-NLS with a nuclear marker in plant cells (Fig. 4B). Moreover, the fusion between full length Avr3 with NLS did not disrupt delivery of the effector to haustoria, and we did not observe the presence of Avr3 in *P. capsici* nuclei within the hyphae (Fig. 4C). However, nuclear localization was observed when the NLS was fused to a truncated version of Avr3 carrying only the first 58 amino acids (the fragment from the SP to EER motif) (Avr3ΔC-Venus-NLS) (Fig. 4D, Supplemental Movie S3).

### Transport of Avr3 towards haustoria depends on its C-terminal region

The localization of Avr3ΔC-Venus-NLS within the nuclei of *P. capsici* led to the question if this effect is caused by (i) the artificial fusion with NLS and competition between secretory and nuclear localization signals or (ii) the absence of the effector C-term fragment that results in impaired secretion. To address this question, we compared the localization pattern of full length Avr3 fused with Venus with its truncated version (Avr3ΔC-Venus) when expressed in *P. capsici*. Whereas Avr3-Venus signal was enriched only around the neck region of haustoria, Avr3ΔC-Venus labeled both haustoria and the hyphae (Fig. 5A and B). At haustoria, the truncated Avr3 version showed two patterns of localization: (i) uniform dispersion inside the haustorium without any accumulation site and (ii) as a ring accumulating at haustorial necks (Fig. 5C). Additionally, the intensity profile confirmed localization of uniformly dispersed Avr3ΔC-Venus signal within the haustorium and the lack of secretion (Fig. 5D).

**Figure 5.**
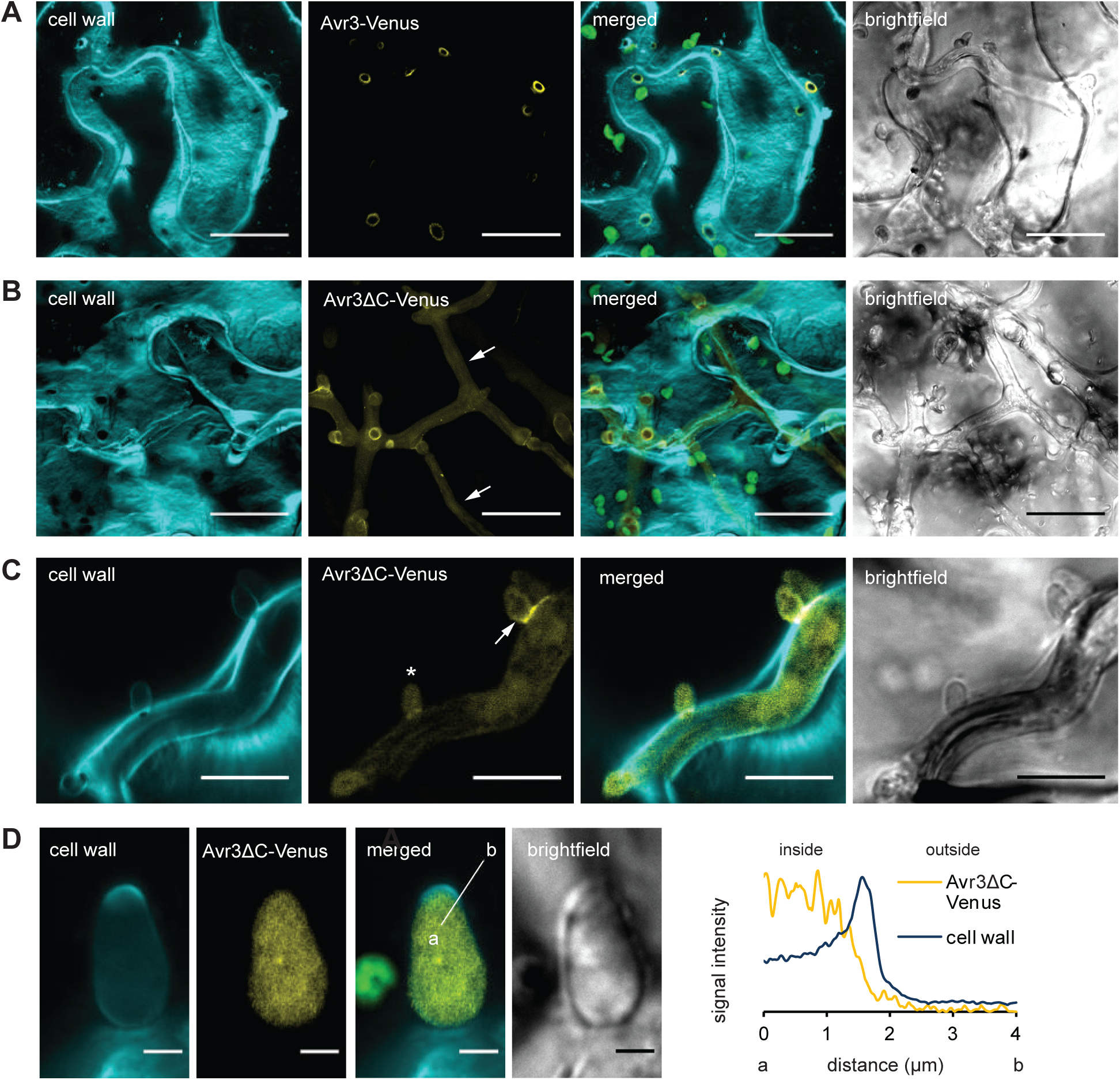
Avr3 C-terminal region is required for secretion from *P.capsici* hyphae. Representative confocal microscopy pictures of *P. capsici* expressing Avr3-Venus (maximal projection z-stack of 8 slices) and (**B**) Avr3 truncated version that does not carry the sequence downstream of EER motif (maximal projection z-stack of 26 slices). Hyphae with retained Avr3ΔC-Venus signal marked with arrows. Scale bars 20 μm. Chloroplasts labeled in green. Avr3ΔC-Venus expressed by *P. capsici* forms two localization patterns at haustoria: an arrow marks haustorium with Avr3ΔC-Venus signal accumulated at the haustorial neck, star marks the haustorium where Avr3ΔC-Venus does not show accumulation site but is dispersed uniformly inside haustorium. Scale bar 10 μm. High magnification of confocal microscopy picture presents haustoria with Avr3ΔC-Venus that is localized inside haustorium and not secreted. Scale bar 2 μm. On the right the relative fluorescence intensity plot of signals detected along the line shown in the merged picture (a-inside, b-outside haustorium). Chloroplasts labeled in green. Cell wall stained with calcofluor white.

Taken together, these data show that the absence of the Avr3 C-terminal part affects Avr3 secretion. A possible explanation is that in the absence of the C-term, the sorting of Avr3 to the secretory pathway is disrupted and as a result, Avr3 remains within the hyphal cytosol. Based on the model proposed by Boevink et al. (2020), the secretion process of RxLR effectors might entail the formation of a structure created by extensions of the endoplasmic reticulum (ER) that subsequently detach and engulf cytoplasm. Interestingly, the effector domain at the C-terminal part was discovered to mediate binding to phosphatidylinositol monophosphates (PIPs) (Yaeno *et al*., 2011) which are known to control membrane trafficking and recruitment of cytosolic proteins (von Blume & Hausser, 2019).

Remarkably, when secreted, the truncated Avr3 (Avr3ΔC-Venus) accumulated at the haustorial neck in the same manner as the full-length Avr3. This outcome is unexpected because the truncated version of Avr3 solely comprises the signal peptide, an RxLR-EER motif fragment and a Venus-tag, lacking any other recognizable effector domain that could determine this specific location. This implies that the RxLR-EER motif alone is sufficient to determine localization at the haustorial neck. Another puzzling aspect is that Avr3 constructs label the interface only at the haustorial neck and not the rest of haustoria body.

### The RxLR-EER motif is sufficient to determine protein localization to the haustorial neck distinct to EHMx

To test the role of the RxLR-EER motif in the localization of the construct to the haustoria neck, we re-examined the accumulation patterns at haustoria in *P. capsici* transformants secreting Avr3-Venus (Fig. 6A), its truncated variant (Avr3ΔC-Venus) (Fig. 6B), and the truncated version tagged with NLS (Avr3ΔC-Venus-NLS) (Fig. 6C). As in previous observations, the truncated Avr3 variants exhibited retention either within the cytosol (Fig. 6B) or the nucleus (Fig. 6C). However, upon reaching the haustoria, both variants displayed localization at the haustorial neck, mirroring the behavior of the full-length Avr3. Notably, localization of these constructs exhibited regional feature without equal continuum distribution in EHMx that surrounds the whole body of haustoria.

**Figure 6.**
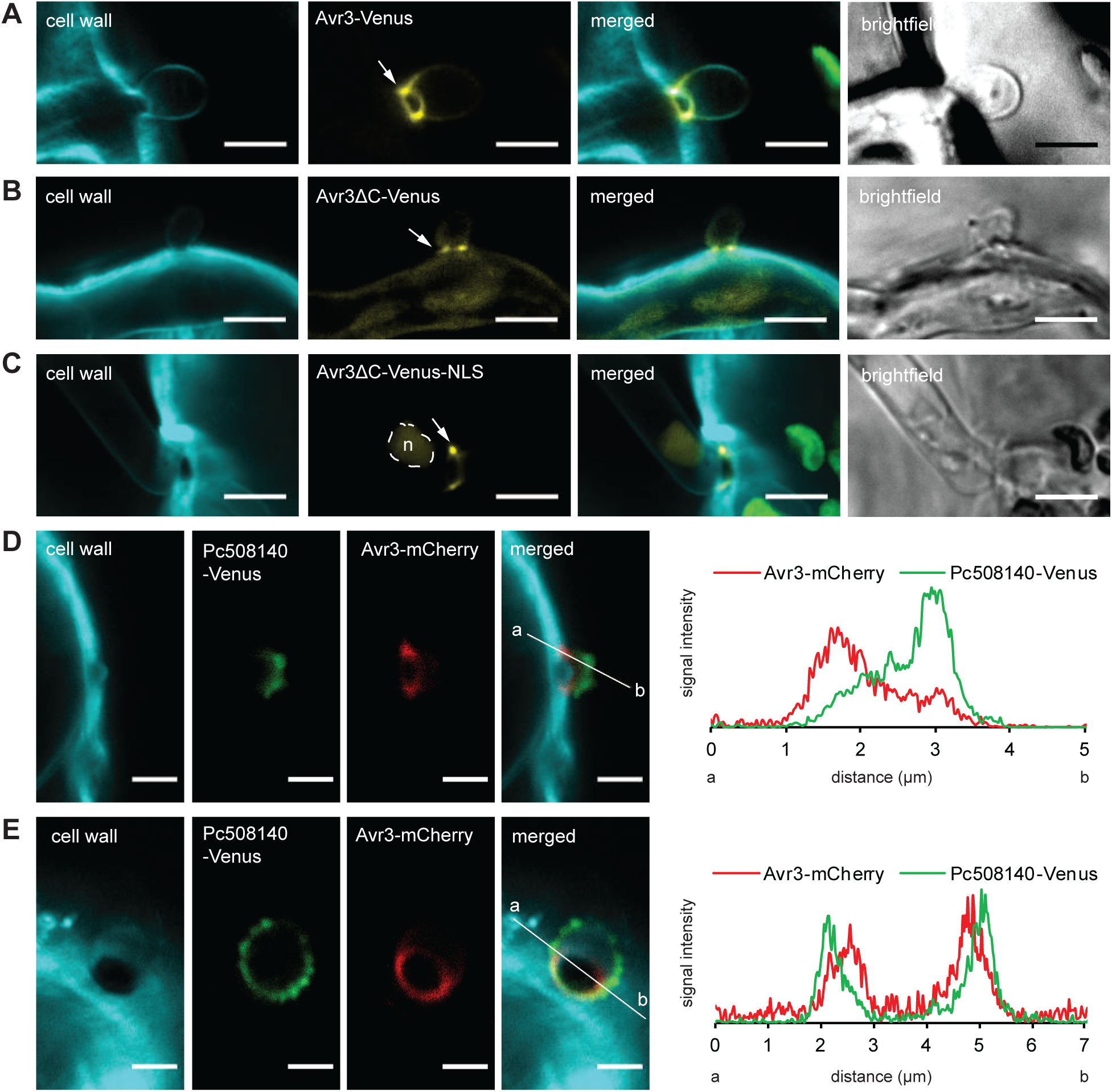
RxLR-EER motif directs Avr3 to the haustorial neck. Representative confocal microscopy pictures of *P. capsici* haustoria with the Venus signal fused with (**A**) Avr3 (**B**) Avr3ΔC and (**C**) Avr3ΔC-NLS. All fusions accumulate at the neck of haustorium (marked with arrows in the pictures **A**-**C** although retention in cytoplasm inside the hypha or the Phytophtora nucleus (n, the shape of the nucleus outlined) (**C**) was also observed. Chloroplasts labeled in green. Scale bars 5 μm. Representative confocal microscopy pictures of Avr3-mCherry and Pc508140-Venus secreted simultaneously from *P. capsici* secretory pocket (**D**) and haustoria (**E**). On the right each panel is accompa nied by relative fluorescence intensity plot of Venus and mCherry signals detected along the line from the point a to b shown in the merged picture. Scale bars 2 μm. Cell wall stained with calcofluor white.

Following this confirmation, we aimed to address two observations: 1.) Why are Avr3 constructs not distributed over the entire EHMx that envelops the whole haustoria body and 2.) Does the localization of Avr3 at haustorial necks depend on the RxLR-EER sequence? One possible explanation for the absence of Avr3 constructs from the EHMx is that the space of EHMx is too limited with restricted protubernaces, whereas the space around the haustorial necks is larger. To test this hypothesis, we investigated the co-localization between Avr3-mCherry and the EHMx marker Pc508140-Venus, which were co-expressed simultaneously in *P. capsici*. The secretory pockets and haustoria exhibited well-developed EHMx with protuberances labeled with Pc508140-Venus signal (Fig. 6D and E). However, in contrast to Pc508140-Venus, Avr3-mCherry preferentially accumulated at the base of the secretory sites and did not disperse around the EHMx. Analysis of the intensity profiles revealed spatial separation between Pc508140-labeled EHMx and the Avr3-positive region at the haustorial neck, indicating that Avr3 resided proximally to EHMx (Fig. 6E). For this reason, we performed a co-localization experiment with Avr3-Venus and the haustorium cell wall. The results showed that in both longitudinal and transverse views, the Avr3-Venus signal co-localized with the haustorial cell wall (Fig. 7A and B).

**Figure 7.**
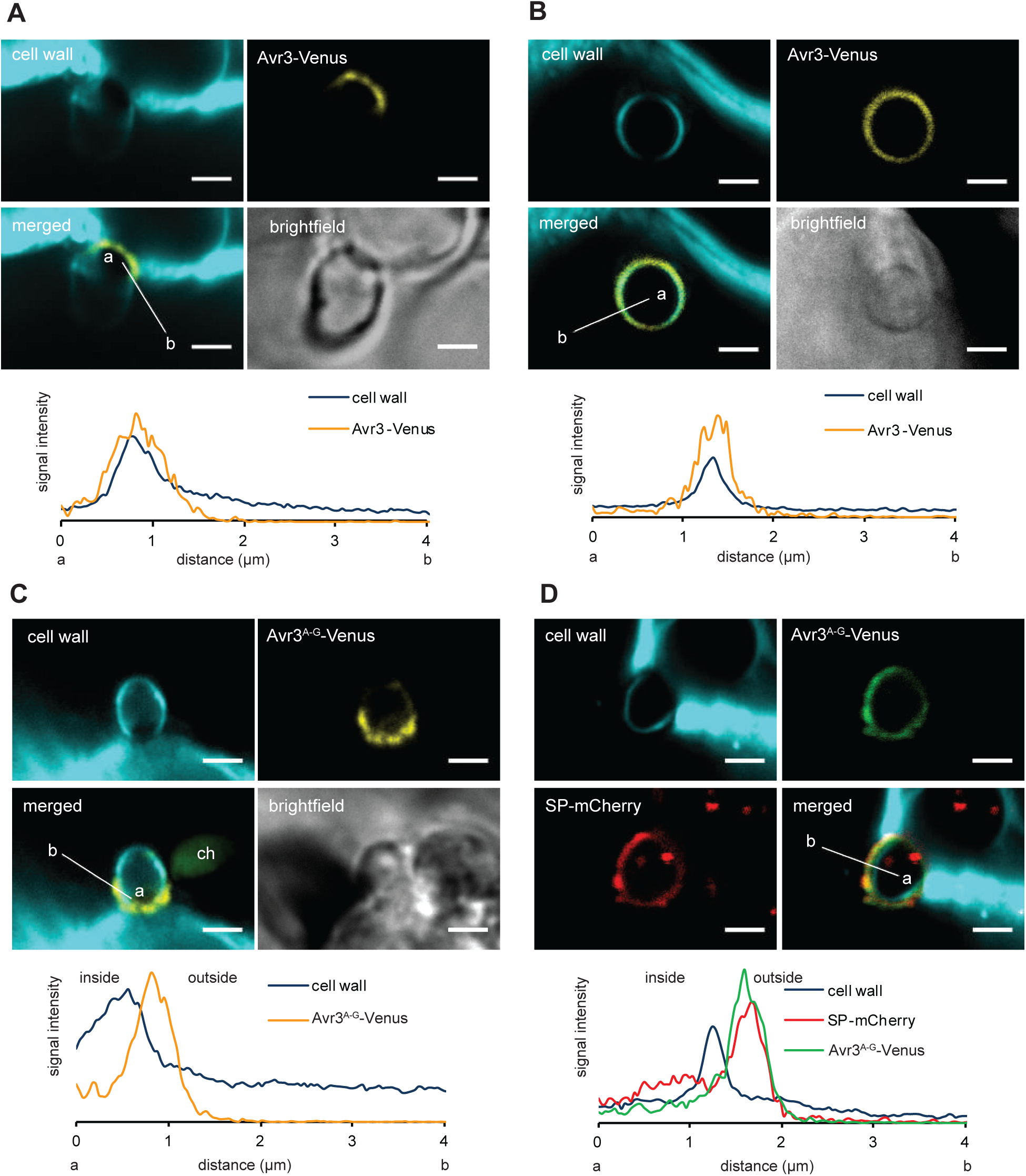
RxLR-EER motif fragment associates Avr3 with the haustoria cell wall whereas its mutation redirects the effector to EHMx protuberances. Representative confocal microscopy pictures of Avr3-Venus secreted by *P. capsici* and colocalizing with haustoria cell wall in (**A**) longitude and (**B**) cross section of haustorium. (**C**) Mutated Avr3 with substituted RxLR-EER motif fragment (Avr3^A-G^-Venus) is secreted by *P. capsici* outside haustorial cell wall and (**D**) colocalizes with secreted free mCherry protein (SP-mCherry) in protuberances of EHMx. On the bottom each panel is accompanied by relative fluorescence intensity plot of signals measured along the line (a-inside, b-outside haustorium) shown in the merged picture. Scale bars 2 μm. Chloroplast labeled in green (ch).

To verify the role of the RxLR-EER motif in Avr3 localization at haustorial necks, we replaced the RxLR motif with alanine and the dxEER sequence with glycine residues. Contrary to Avr3-Venus, the mutant with alanine-glycine substitutions (Avr3^A-G^-Venus) did not accumulate at haustorial necks but displayed localization within the EHMx protuberances (Fig. 7C, Supplemental Fig. S7). This finding was further confirmed by co-localization of mutated Avr3-Venus with SP-mCherry as an EHMx marker. Both signals were found to co-localize outside the haustorial cell wall, providing evidence for their secretion into the EHMx (Fig. 7D).

Taken together, these experiments reveal that prior to translocation into the plant cell, Avr3 is not targeted to the EHMx like other apoplastic proteins. Instead, it preferentially accumulates in a distinct site at the neck region of the secretory sites. Furthermore, the co-localization with the haustorial cell wall suggests that the Avr3 resides within the layer of the haustorial cell wall/plasma membrane. Thus, our findings unequivocally demonstrate that Avr3 loses association with the haustorial neck region and undergoes re-localization to the EHMx when the RxLR-EER motif is mutated. This raises an intriguing question regarding how the RxLR-EER motif influences the connection of Avr3 with the haustorial neck. Until now, it has been commonly assumed that the RxLR-motif undergoes cleavage prior to, or during, effector secretion (Wawra *et al*., 2017; Wang *et al*., 2023).

### The RxLR motif remains an integral part of Avr3 effector during *P. capsici* infection in plant tissue

According to Wawra *et al*. (2017), the Avr3 effector secreted to the media by axenically cultured *P. infestans* lacks the RxLR motif. This observation led to the conclusion that the RxLR motif cannot be directly implicated in effector uptake by the host cell, but it must function prior to secretion, possibly during recruitment to the appropriate secretory pathway. We thus investigated the relation between RxLR motif cleavage and Avr3 association with the haustorial neck region. We created *P. capsici* transformants that express double tagged versions of Avr3: with mCherry inserted between the SP and the RxLR motif of Avr3, and a Venus-tag positioned at the C-terminal end of Avr3. The rationale behind this strategy was that the separation of the Venus and mCherry signals will unveil the compartment in which the cleavage takes place: while the C-terminal Venus-tag will label the core structure of Avr3, the mCherry signal will indicate the localization of the cleaved RxLR motif. In comparison to the native effectors, the obtained construct has an abnormally large size. To control for this, we developed a similar Avr3 chimera where the RxLR-EER motif fragment was substituted with alanine-glycine residues (mCherry-Avr3^A-G^-Venus-FLAG).

Both chimeras were targeted by *P. capsici* towards the haustoria. However, the mCherry-Avr3-Venus-FLAG chimera exhibited distinct localization patterns of fluorescent tags at the haustorial sites: mCherry appeared distributed within the EHMx protuberances, whereas Venus accumulated at the neck of the haustorium, consistent with the localization we demonstrated for Avr3. The separation of mCherry and Venus signals was validated by intensity profile (Fig. 8A). In contrast, the Avr3 chimera with a mutated RxLR-EER motif fragment did not display any signal separation at the haustorium (Fig. 8B). The detection of both mCherry and Venus signals at the haustorium strongly indicates that the Avr3 chimeras reach the haustorium in an intact form. However, the separation of fluorescent tags within the haustorial region, observed via confocal microscopy for Avr3 but not its mutated version, implies cleavage occurring within the Avr3 protein sequence that comprises the RxLR-EER motif.

**Figure 8.**
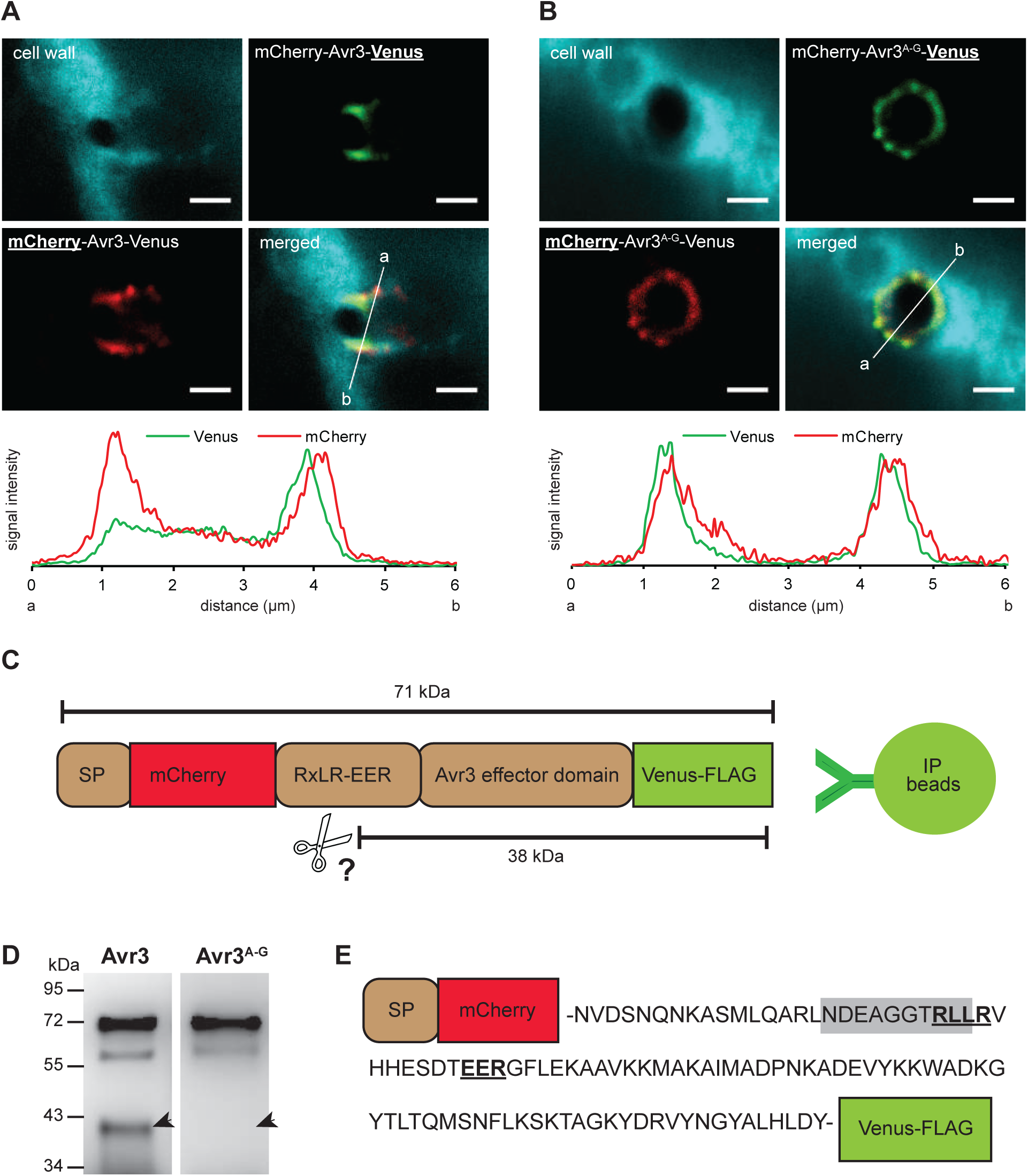
Cleavage site in double tagged Avr3 secreted by *P. capsci* during plant infection is dependent on RxLR-EER motif fragment but the motif itself remains intact. Representative confocal microscopy pictures of *P. capsici* haustoria secreting mCherry and Venus double tagged (**A**) Avr3 and (**B**) Avr3 mutated version where RxLR-EER was substituted by alanine and glycine residues. The channel observed for each tag underlined in the construct name, scale bars 2 μm. Bottom panels show relative fluorescence intensity plots of both mCherry and Venus signals detected along the arrow shown in the merged picture. (**C**) Schematic draft of the cleavage position and sizes of proteins released after anticipated cleavage of RxLR motif when Avr3 is immunoprecipitated on its C-terminal Venus tag. (**D**) Immunoblot of double tagged Avr3 and its mutant lacking RxLR-EER motif (Avr3^A-G^) immunoprecipitated with GFP antibody from *N. benthamiana* tissue infected with *P. capsici* transformants. Protein fragment of approximate 38kDa size expected to be released after cleavage present in Avr3 sample but absent in Avr3^A-G^ marked with arrows. This experiment was conducted twice on independent batches of infected plant tissue, with similar results. (**E**) Schematic representation of *P. capsici* Avr3 aminoacid sequence tagged with mCherry and Venus. Protein fragment of 38kDa size cleaved from Avr3 contains RxLR motif as detected by LC-MS. The peptide detected by LC-MS carrying RxLR motif highlighted in gray. The RxLR and EER motives are underlined.

To further verify this observation, we analyzed bands derived from Avr3 chimera constructs by immunoblot. *P. capsici* transformants that overexpressed Avr3 chimeras (wild type Avr3 and with mutated RxLR-EER motif, respectively) were grown on *N. benthamiana* tissue. Next, the infected tissue was used to immunoprecipitate Avr3 chimeras using an antibody directed against Venus. For both chimeras, we identified bands at 71kDa, consistent with the anticipated size for intact mCherry-Avr3-Venus-FLAG proteins (Fig. 8C). However, the presence of an additional 38kDa band, matching the size of cleaved Avr3-Venus, was exclusively observed in the immunoblot for the Avr3 containing the RxLR-EER fragment (Fig. 8D).

To confirm the nature of the 38kDa protein specific for the wild-type Avr3 chimera and to identify its cleavage site, we isolated the bands for analysis via mass spectrometry (MS). In a previous study with *P. infestans* Avr3, the authors employed trypsin-digested samples in their MS study and identified a cleavage site between the RxLR and EER motif (Wawra *et al*., 2017). Since trypsin specifically cleaves peptide bonds at lysine and arginine residues (Olsen *et al*., 2004), we aimed to avoid the potential proteolysis of the RxLR motif into fragments that might be undetectable. Therefore, to retain the RxLR sequence within the pool of analyzable peptides, we used elastase, which exhibits a less specific cleavage pattern known to preferentially target the carboxyl-terminal side of small neutral amino acids (Wang *et al*., 2008; Meyer *et al*., 2014). The MS analysis of the 38kDa protein revealed the presence of a peptide containing the RxLR motif, pinpointing the cleavage site between the mCherry fusion and the RxLR motif in the double-tagged Avr3 (Fig. 8E, Supplemental Dataset S1). This finding shows that during infection, the RxLR-EER motif remains intact and is not cleaved off from the secreted *P. capsici* effector Avr3.

## DISCUSSION

Effectors serve as pivotal molecules that increase the potential of pathogens to infect and sustain themselves in their hosts, hence they are also referred to as virulence factors. In recent years, bioinformatic predictions combined with transcriptome and secretome analyses across diverse Phytophthora species, have revealed the proteins that are potential virulence factors (Raffaele *et al*., 2010; Meijer *et al*., 2014; Zuluaga *et al*., 2016; Evangelisti *et al*., 2017). There has been also a significant progress in understanding their mechanisms of action and the role of effectors for plant host invasion and suppression of defenses (Tomczynska *et al*., 2018; Judelson & Ah-Fong, 2019; Tomczynska *et al*., 2020). Little is known, however, about the secretion of Phytophthora effectors. There is also considerable controversy surrounding the transport mechanism of RxLR effectors that are translocated to the plant symplast (Dou *et al*., 2008; Kale *et al*., 2010; Wawra *et al*., 2013; Petre *et al*., 2016).

The motivation for our study was to systematically address the functional role of the RxLR motif in effector transport, secretion, and subcellular localization using oomycete effectors expressed in *P. capsici*. During infection, the Phytophtora transcriptome and proteome undergo massive reprogramming that is coordinated by the interaction with the plant host (Savidor *et al*., 2008; Jupe *et al*., 2013; Wang, S *et al*., 2021). Changes occur also at the level of protein secretion, as described for *P. plurivora* (Severino *et al*., 2014). After infection, the haustoria, structures formed by Phytophthora during the biotrophic phase, are sites of default protein secretion where various proteins with a signal peptide are directed (Kagda *et al*., 2020). Haustoria were shown to serve a crucial function in the secretion of both apoplastic and cytosolic virulence factors (Wang *et al*., 2017; Wang *et al*., 2018).

By employing *P. capsici* transformants that secrete mCherry fused to the signal peptide (SP-mCherry), we showed that mCherry accumulated outside haustoria in EHMx, consistent with previous findings of Kagda et al. (2020). However, within the same transformants, the hyphae that failed to penetrate the host tissue did not exhibit any evident SP-mCherry secretion, illustrating the pivotal role played by the haustorial interface formed by both the pathogen and the plant during infection and its importance in investigating biologically relevant aspects of effector transport. This is particularly crucial in studying RxLR effector delivery, as the interface can significantly influence, if not directly facilitate, the translocation of these effectors (Petre & Kamoun, 2014).

The study of Phytophthora secretory processes has been seriously constrained by the demanding confocal microscopy experiments, which are time-consuming and require specific skills and experience (Boevink *et al*., 2020). In addition, Phytophthora transformation itself is a challenging procedure with low efficiency (Ghimire *et al*., 2022). We therefore employed an approach described by Fang et al. (2017) using transiently transformed *P. capsici* and an infection protocol with agar plugs that provides uniform penetration and a robust system for infection (Taylor & Grünwald, 2021). Furthermore, we reasoned that the Phytophthora cytosol does not provide a good subcellular marker for haustoria organization. Therefore, we successfully employed fluorescent dyes to stain callose and cellulose to label the *P. capsici* cell wall, which offers a straightforward and reliable strategy for visualizing the interface at Phytophthora haustoria.

A striking finding from our study is the demonstration that the intracellular RxLR effector Avr3, which is delivered to the plant cell, shows a ring-shaped localization at the haustorial neck region and not the rest of the EHMx around the haustorial body. Further, the RxLR-EER motif is responsible for Avr3 association with the haustorial neck region and in contrast to the previous report (Wawra *et al*., 2017), this motif is not cleaved off but remains an integral part of Avr3 after secretion. The correlation between the facts that (i) effector translocation requires presence of haustoria and RxLR-EER motif (Whisson *et al*., 2007) and (ii) RxLR-EER motif mediates targeting of the protein to the haustorial neck support our hypothesis that this specific localization is linked to the mechanism of effector translocation. The localization of *P. capsici* RxLR effector Avr3 at the haustorial neck region aligns with previous observations that RxLR effectors accumulate at the haustorial neck when secreted by transformants of *P. parasitica* (Huang *et al*., 2019) and *P. infestans* (Boevink *et al*., 2020). Moreover, a similar characteristic ring at the neck region of biotrophic primary hyphae was described for *Colletotrichum* species (Irieda *et al*., 2014; Irieda *et al*., 2016; Ogawa *et al*., 2021). A ring-like fluorescence structure is the site of focal accumulation for mCherry-tagged intracellular effectors DN3 and NIS1. In contrast to DN3-mCherry, mCherry fused to the signal peptide of DN3 (SP-mCherry) did not form a ring signal efficiently. The authors also concluded that the region at the pathogen-plant interface where DN3 and NIS1 localize comprises membranous components. Notably, it is formed only in the presence of the host plant but not on an artificial, penetrable substratum (Irieda *et al*., 2014). There is evidence that the plant-pathogen interface at haustorial necks differs from the one at the haustorial body. In contrast to the distal end of the haustorium, the area around the base displayed a distinct structure, appearing as a thin layer of highly electron-dense material (Mims *et al*., 2004).

In our study, the Avr3-positive haustorial neck region of Phytophthora haustoria is distinct from the region of the EHMx where free proteins are secreted. A plausible interpretation of the separation between the neck region and the rest of EHMx is that the neck region is an environment that is isolated from the rest of EHMx or alternatively, possesses a distinct structural composition. Within the wall of the haustorial neck, rust fungi were shown to form a structure termed the neckband. This noticeable thickened region seals the fungal plasma membrane with the invaginated host-cell plasma membrane. It acts as a permeability barrier and is believed to create a sealed subcompartment by preventing outflow of material from the EHMx to the plant apoplast (Heath, 1976; Mims *et al*., 2004). While neckbands have been also reported for powdery mildews, it seems that these structures are absent in haustoria-forming oomycetes (Perfect & Green, 2001). Apart from an exceptional case of *Albugo candida*, which forms neckband-like structures that differ from those formed by fungi (Woods & Gay, 1983; Soylu, 2004), neckbands have not been documented for Phytophthora species (Shimony & Friend, 1975; Hohl & Stössel, 1976; Hohl & Suter, 1976). In fact, it rather seems that the *P. infestans* EHMx is continuous with the plant apoplast (Whisson *et al*., 2007).

We observed that the Avr3-positive region at haustorial necks exhibited co-localization with the Phytophthora cell wall, similar to the plasma membrane marker. The membranous origin of Avr3 positive structures at the haustorial neck in our study can be a plausible explanation why this haustorium region is distinct and physically separated from the EHMx where free proteins are released. Before, it was hypothesized that translocation of *P. infestans* Avr3a requires not only the presence of the haustorium and extrahaustorial matrix but also a mechanism that would facilitate the passage of RxLR effectors to the plant cell (Whisson *et al*., 2007). Later, it was also concluded that free effector proteins applied in the plant apoplast do not have ability to enter host cells (Petre *et al*., 2016). Moreover, even during Phytophthora infection, RxLR effectors that are expressed and secreted by the plant cell into the apoplast do not re-enter the plant cell. Instead, they remain in the apoplast surrounding haustoria (Wang *et al*., 2017). In several fungal species, the extrahaustorial matrix around the haustorial neck contains tubular elements (Manners & Gay, 1983; Mims *et al*., 2003; O’Connell & Panstruga, 2006). Although there is no evidence that they transport substances across the interface, the authors suggest they may secrete fungal effectors (Roth *et al*., 2019). A similar hypothesis proposed that these membrane connections might help pathogens access the host endomembrane system, potentially serving as pathways for protein trafficking into host cells (Catanzariti *et al*., 2007).

Given that the RxLR-EER motif determines the ability of proteins to be delivered through the plant cell membrane (Whisson *et al*., 2007), and since in our studies, the RxLR-EER motif was sufficient to guide the Venus tag to the ring-shaped structure at the haustorial neck and mimic localization of full length Avr3, it is tempting to consider it as an interface formed by the pathogen to facilitate effector translocation. The best-investigated machinery for effector translocation in plants is the biotrophic interfacial complex (BIC) of the rice blast fungus *M. oryzae*. It is a membranous cap formed by the extra-invasive hyphal membrane and its localization is independent from extracellular space outside the invasive hyphae, where apoplastic effectors reside. Fluorescently labelled intracellular effectors are routinely observed in the BIC before they can be detected in the plant cell nucleus (Kankanala *et al*., 2007; Khang *et al*., 2010). More detailed studies showed that within the BIC, cytoplasmic effectors are packaged in dynamic vesicle-like membranous effector compartments (MECs) and are taken up by plants via endocytosis (Oliveira-Garcia *et al*., 2023). Likewise, in endosome fractionation experiments, *P. infestans* RXLR effector Pi04314 was found to be co-enriched with membrane-associated proteins originating from plant clathrin-coated endosomes (Wang *et al*., 2023). In the light of these findings, we propose that the region at the haustorial neck where Avr3 accumulates is a structure that is functionally analogous to the BIC of *M. oryzae*.

In previous experiments with the native, non-tagged AVR3a secreted by an axenic *P. infestans* culture into the media, mass spectrometry (MS) analysis did not detect the RxLR motif. Furthermore, it was concluded that the RxLR motif is cleaved off, which appears in a way similar to the fate of the PEXEL motif of *P. falciparum* effectors. However, neither the plasmepsin V protease of *P. falciparum* that cleaves the PEXEL motif, nor other 10 investigated aspartic protease showed proteolytic activity towards RxLR proteins (Wawra *et al*., 2017). We provide evidence that the RxLR-EER motif stays intact after Avr3 effector secretion from *P. capsici* haustoria during infection. Earlier research demonstrated that both variants of Avr3a, whether with or without the RXLR-EER fragment, are targeted to haustoria, indicating that RxLR-EER is not necessary for secretion. The difference in peri-haustorial localization of Avr3 lacking RxLR-EER was interpreted as a consequence of the impaired translocation to the plant cell, resulting in effector accumulation in the EHMx (Whisson *et al*., 2007). Similarly, intense fluorescence signal confined to the haustorial base observed for RxLR effector Pi04314-mRFP was interpreted as a region of high level of effector secretion (Boevink *et al*., 2020). Given that in our experiments the RxLR-EER motif is not cleaved, we postulate that the RxLR-EER motif mediates protein targeting to the haustorial neck region, which further facilitates translocation to the host cell. Accordingly, mutated Avr3 version entered the EHMx space instead of being directed to the haustorial neck.

While we demonstrate the association of RxLR-EER proteins with the neck at haustoria, the exact mechanism of this association remains unknown. It remains uncertain whether the RxLR-EER motif provides a structural basis for the effector protein or facilitates direct attachment. Wawra and colleagues (2017) suggested that the RxLR-EER motif serves as a sorting signal to the effector secretory pathway, whereas Kale and colleagues (2010) proposed that RxLR-EER binds to PI3Ps. However, according to Yaeno *et al*. (2011), PI3P3 binding is attributed to the C-terminal region of the effector. Possibly, the correct sorting into the secretory pathway is linked with the capability for translocation; thus, the RxLR-EER motif could be a sorting signal that attaches Avr3 to the surface membrane of secretory vesicles and creates further attachment at the haustorial neck. Future studies should investigate whether accumulation of RxLR effectors at the haustorial neck is a common phenomenon in Phytophthora species and address its origin and functional relevance in the translocation mechanism (Supplemental Fig. S8).

## MATERIAL AND METHODS

### Plasmid construction

Gene-specific primers with in-phusion overhangs were designed with use of Vazyme primer design tool for Multi-Fragment Assembly. Genes were amplified from genomic DNA of *P.capsici* isolate Pc263 and Avh110 was amplified from *P. sojae* strain P6497. *P. capsici* plasma membrane marker (PM-mCherry) was generated containing the myristoylation/ palmitoylation sequence MGCGCSSHPED described elsewhere (Rozbicki *et al*., 2015). Sequence of the nuclear localization signal from simian virus large T-antigen (three tandem repeats of PKKKRKV) (Khang *et al*., 2010) were fused with Avr3 for its targeting to the plant nucleus (Avr3-Venus-NLS). Detailed sequences of the primers used in this study and the final phusion proteins are listed in Supplemental Table S1. PCR amplifications were conducted using Phusion® high-fidelity DNA polymerase (New England Biolabs, Ipswich, MA, USA). PCR products were purified from the gel and cloned into the backbone of Phytophthora expression vector pMCherryN (Ah-Fong & Judelson, 2011) between *Nhe*I and *Sac*II restriction sites with Vazyme ClonExpress Ultra One Step Cloning Kit according to the manufacturer’s protocol. Resulting plasmids were transformed and propagated in DH5-Alpha *E.coli* grown on LB (Luria-Bretani broth medium) with 100 μg/mL ampicillin as the selection marker. The expression of the resulting fusion construct was driven under the control of the promoter and terminator from the *Ham34* gene.

### Phytophthora transformation

*P. capsici* wild type isolate Pc263 was routinely grown in cleared V8 medium at 25 °C in the dark. The approach of Phytophthora transient transformation was adopted from (Fang & Tyler, 2016; Fang *et al*., 2017). Plasmids for transformation were isolated with PureLink HiPure Plasmid Maxiprep Kit (Invitrogen). Transformation procedure was performed using a modified polyethylene glycol protocol (Fang & Tyler, 2016). Protoplasting step was done with 0.32% Lysing Enzymes from *Trichoderma harzianum* (L1412, Sigma) and 0.25% Cellulase from *Trichoderma reesei* (C8546, Sigma). *P.capsici* independent transient transformants (mycelia that recovered spatially separated after selection) were maintained on V8 medium supplemented with 50 μg/mL G418 (Geneticin, AG Scientific, San Diego, California, USA). They were analyzed within 3 weeks after their recovery in selection media. First, they were used to infect *N. benthamiana* leaves and pre-selected visually for the presence of Venus and/or mCherry under confocal laser scanning microscope. As positive transformants showed the same fluorescence patterns with varying intensities, only those with strongest fluorescent signal were further maintained. From each batch of transformants, at least six independent representative mycelia that exhibited normal, radius-shaped growth on the media and high number of hyphae with the fluorescent signal across the entire infection side were maintained for further microscopy observation. The expression of the investigated construct was further confirmed in two independent transformants by detecting proteins immunoprecipitated from *N. benthamiana* tissue infected with these transformants (Supplemental Fig. S8).

### *Nicotiana benthamiana* infection

*N. benthamiana* plants were grown in standard conditions (Tomczynska *et al*., 2018) and two middle leaves from four week old plants were used for infection. Infection was performed on detached leaves inoculated with a single mycelial plug (Taylor & Grünwald, 2021). For each *P.capsici* transformant, a 6mm^2^ plug cut with a sharp blade from the external edge of 3 day old mycelium was placed on abaxial side of *N. benthamiana* wounded leaves and incubated in the darkness at 25°C in high humidity. Leaf pieces were analyzed 18h post-infection for the localization of the fluorescent protein fusions in *P. capsici* transformants and as tissue for immunoprecipitation.

### Staining procedures

Staining with fluorescent dyes was performed by syringe infiltration of working solutions through the abaxial surface of *N. benthamiana* leaves infected with *P. capsici* transformants and observed immediately. Cell wall was stained with aniline blue or calcofluor white 0.1% (w/v, in 0,1M sodium phosphate pH 9.0 and water, respectively), plant nuclei were stained with 4’,6-diamidino-2-phenylindole (DAPI) solution (100 ng/ml) as described before (Jiang *et al*., 2021).

### Confocal microscopy

*Microscopy was* performed on a confocal laser scanning microscope STELLARIS 8 FALCON using 40x apochromat water immersion objective. Excitation/emission wavelengths were 405 nm/420-490 nm for DAPI, aniline blue and calcofluor white, 514nm/520-560 nm for Venus, 561 nm/595-670 nm for mCherry, 561nm/730-800 nm for visualization of chloroplasts. Images were taken in the lesion area with minimal hyphal colonization to reduce the potential risk of interference from autofluorescence due to infection-related cell damage. The images shown are representative of over 100 haustoria, observed from at least three independent biological replicates. Images were projected and processed with use of ImageJ software (https://imagej.net).

### Immunodetection of tagged constructs

*P. capsici* proteins were immunoprecipitated from 5g of *N. benthamiana* leaf tissue that was infected by respective transformants expressing tagged proteins. Protein extraction was carried out with 25mM HEPES, 150mM NaCl, 0.9% Triton X-100, 1mM EDTA and supplemented with cOmplete protease inhibitor coctail tablets (Roche) (Tomczynska *et al*., 2018). For immunoprecipitation of PM-mCherry, a combination of 0.4% Triton X-100 with 0.03% SDS was used. Extracted proteins were incubated with 18ul of beads slurry for 2h at 4°C on the rotation wheel. To immunoprecipitate Venus and mCherry fusion proteins, respectively RFP- and GFP-Trap Agarose (Chromotek) were used. Beads were washed three times with the extraction buffer, resuspended with 30μl of 2.5x SDS-PAGE sample-loading buffer and 15μl of the sample was loaded on the gel. SDS– PAGE and immunoblotting were performed with standard procedures. Detection of Venus-tagged proteins was based on anti-GFP tag Polyclonal IgG antibody (Chromotek) (1:1000 dilution) and anti-Rabbit IgG conjugated with HRP (Agisera) (1:10 000 dilution). For detection of mCherry-tagged proteins monoclonal igG2c RFP antibody (Chromotek) (1:2000 dilution) and anti-mouse IgG conjugated with HRP (Jakson Immunoreserach) (1:2000 dilution) were used.

### Mass Spectrometry

Twenty five g of *N. benthamiana* tissue infected with *P. capsici* transformants expressing SP-mCherry-Avr3-Venus and its version with mutated RxLR-EER fragment (SP-mCherry-Avr3^A-G^-Venus) respectively was used to immunoprecipite Venus tagged constructs. For each sample, immunoprecipitated proteins were denatured, reduced, alkylated and separated on gel. Next, five equal-sized gel slices were excised and digested with elastase. The resulting peptide mixtures were processed on STAGE tips (Shevchenko *et al*., 2006; Rappsilber *et al*., 2007). LC-MS/MS measurements were performed on the Exploris 480 mass spectrometer coupled to an EasyLC 1200 nanoflow-HPLC (Thermo Scientific). Peptides were separated on a fused silica HPLC-column tip (I.D. 75 μm, New Objective, self-packed with ReproSil-Pur 120 C18-AQ, 1.9 μm to a length of 20 cm) using a gradient of A (0.1% formic acid in water) and B (0.1% formic acid in 80% acetonitrile in water). The samples were loaded with 0% B with a flow rate of 600 nL/min; peptides were separated by 5%–30% B within 85 min with a flow rate of 250 nL/min. Spray voltage was set to 2.3 kV and the ion-transfer tube temperature to 250°C with no sheath and auxiliary gas. Mass spectrometers were operated in the data-dependent mode. After each MS scan (mass range m/z = 370 – 1750; resolution: 120’000) a maximum of twenty MS/MS scans were performed using an isolation window of 1.3, a normalized collision energy of 28%, a target AGC of 50% and a resolution of 15’000. MS raw files were analyzed using MaxQuant software (Cox & Mann, 2008) using the whole sequence of fusion protein and common contaminants was used as a reference. Carbamidomethylcysteine was set as fixed modification and protein amino-terminal acetylation and oxidation of methionine were set as variable modifications. The MS/MS tolerance was set to 20 ppm. Search was done with unspecific digestion mode. Peptide and protein FDR based on a forward-reverse database were set to 0.01, minimum peptide length was set to 7, and minimal number of peptides for protein identification was set to one, which must be unique. The “match-between-run” option was used with a time window of 0.7 min.

### Accession numbers

*Pc508140 (pending), NLP (pending), elicitin* (JX948084.1), *XEG1* (*pending*), *Avr3* (*pending*), *Avh110* (JN253868.1), *RxLR7* (KM068099), *FiB* (KY452016).

## AUTHORS’ CONTRIBUTIONS

IT, MS, DR and MG conceived the research plans. IT conducted experimental work. MS performed mass-spectrometry analysis. IT wrote the article with contributions from all authors. All authors red and approved the final manuscript.

## ACKNOWLEDGEMENTS

This work is dedicated to the memory of Prof. Felix Mauch. The authors would like to thank Prof. Laure Weisskopf, Prof. Stefanie Ranf-Zipproth and Prof. Philippe Reymond for providing support. The project was initiated through an Early Postdoc.Mobility SNSF grant 188054 to IT at Oregon State University, USA and supported by the Swiss National Science Foundation grant 310030_197563 to MMG. IT would like to acknowledge Prof. Brett Tyler and members of Sequencing and Confocal Microscopy Facilities of the Center for Genome Research and Biocomputing at Oregon State University, USA for valuable guidance and critical reviews of the project.

---

**Supplemental Figure S1.**
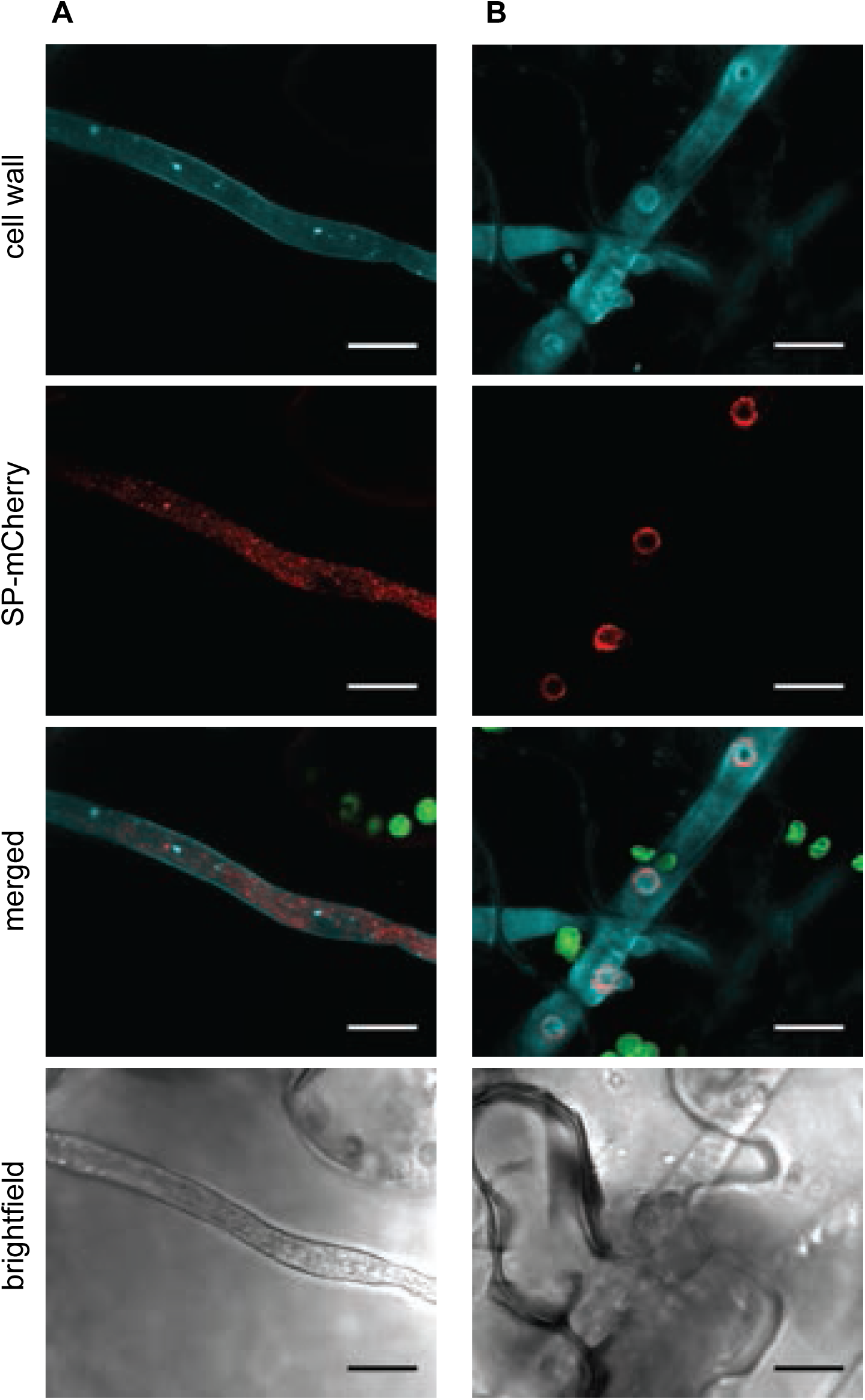
Secretory mCherry marker shows accumulation inside non-penetrating *P. capsici* hyphae and at haustoria penetrating the host cells. Confocal microscopy pictures of *P. capsici* expressing mCherry fused with signal peptide (SP-mCherry). (**A**) Hyphae growing on the surface of *N. benthamiana* epidermis without formation of haustoria, maximal projection z-stack of 11 slices. (**B**) Hyphae penetrating epidermal cells, maximal projection z-stack of 8 slices. Scale bars 10 μm, cell wall stained with aniline blue, chloroplasts labeled in green.

**Supplemental Figure S2.**
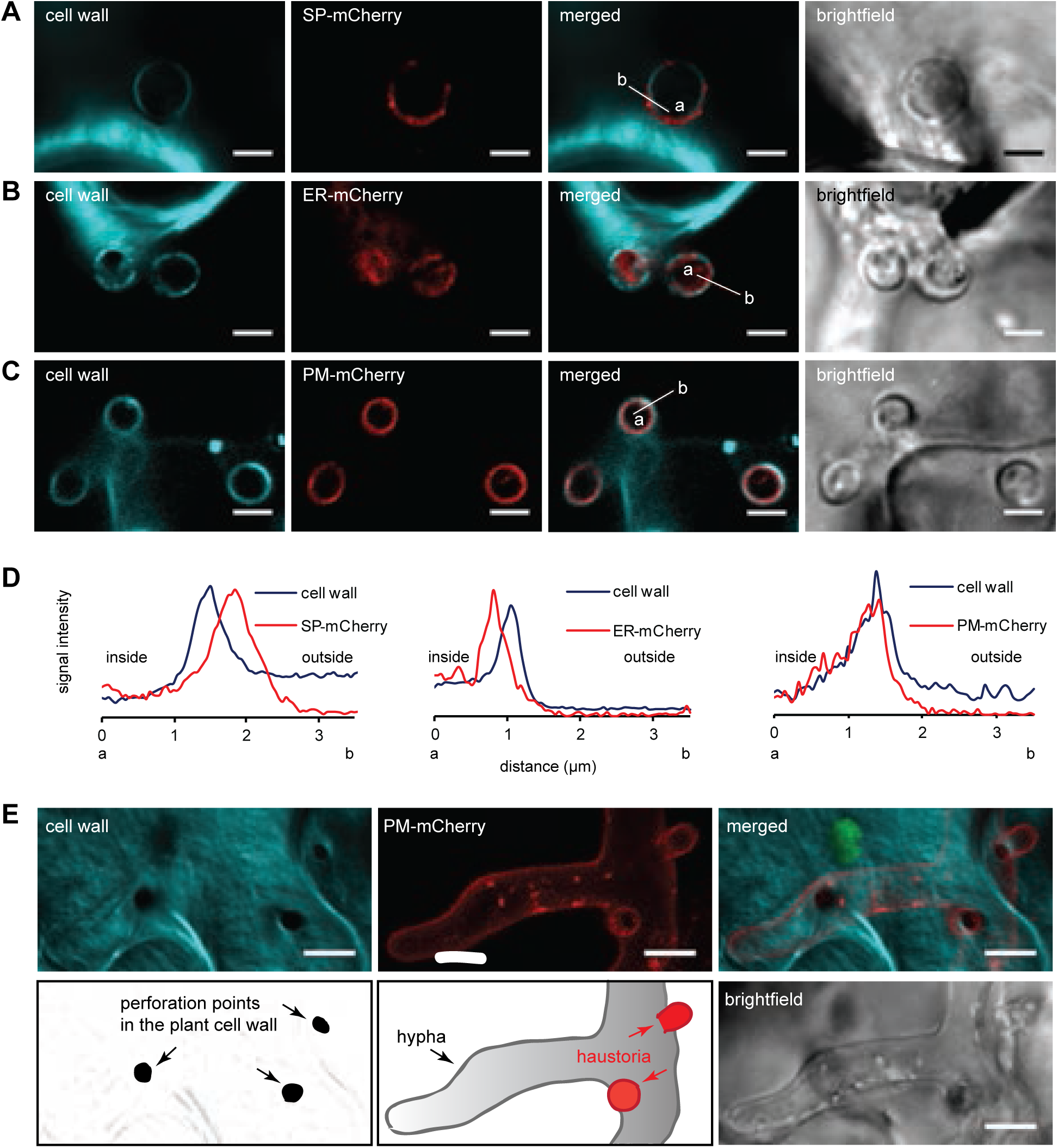
Staining of the cell wall perimeter highlights the interface surrounding *P. capsici* haustoria during infection in *N. benthamiana* epidermal cells. Representative confocal microscopy pictures of co-localization between cell wall stained with fluorescent dyes and *P. capsici* expressing mCherry construct that is either (**A**) secreted (signal peptide fusion), (**B**) retained in endoplasmic reticulum (KDEL fusion) or (**C**) directed to the plasma membrane (myristylation signal), scale bar 3 μm. (**D**) Relative fluorescence intensity plots of cell wall and mCherry signals detected along the line (a-inside, b-outside haustorium) depicted in merged images for constructs shown in A-C. (**E**) Perforation points in the plant cell wall can be formed also in the absence of haustoria. Confocal microscopy z-stack picture (maxi mal projection z-stack of 13 slices) of *P. capsici* with mCherry labelled plasma membrane penetrating the host cell, scale bar 5 μm. Chloroplasts labeled in green. Calcofluor white used for cell wall staining in A and B, aniline blue for C and E.

**Supplemental Figure S3.**
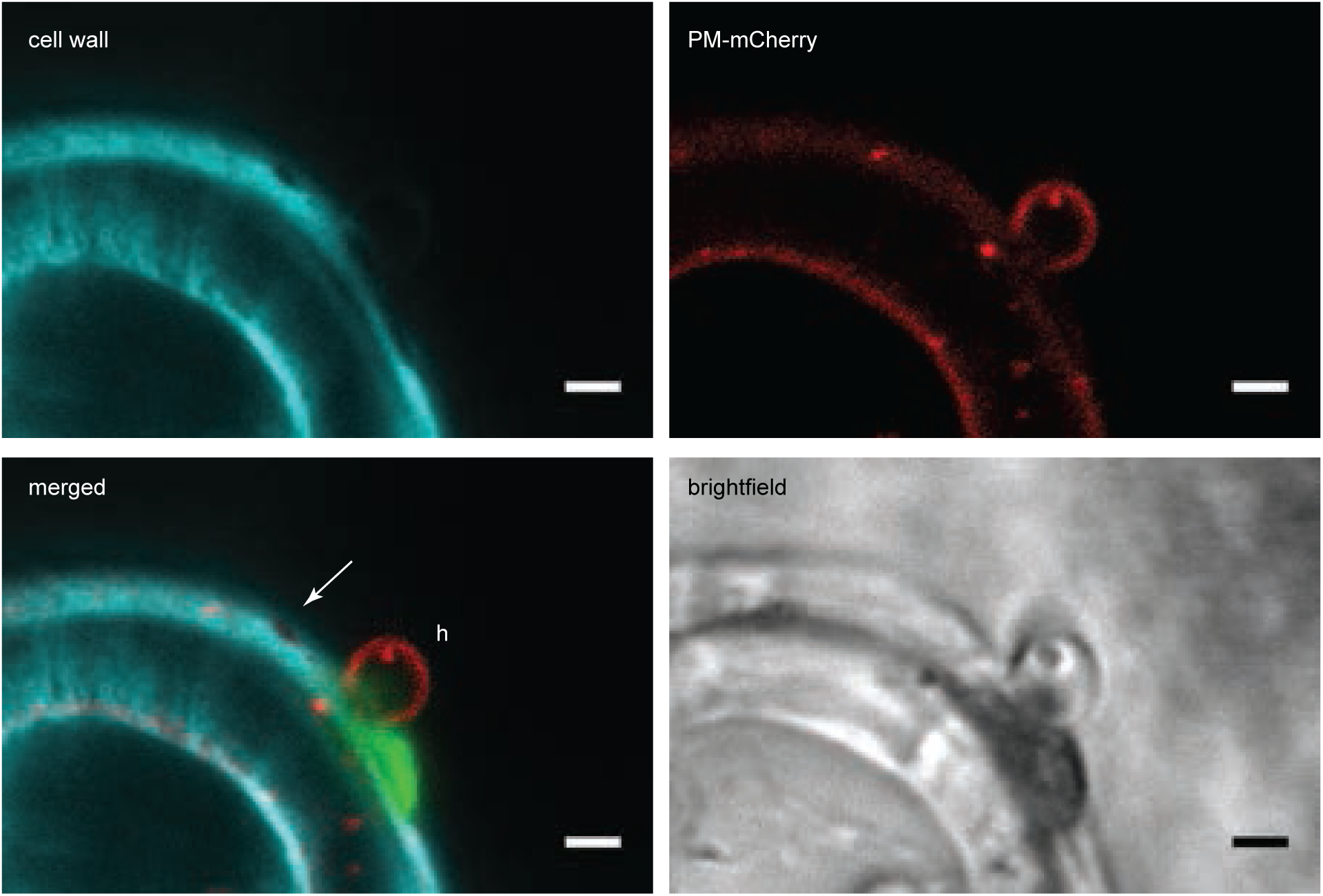
Perforation points on *P. capsici*-*N. benthamiana* cell wall interface appear independently from haustoria. Representative confocal microscopy picture shows side by side compari son between haustorial and haustorial-less perforation point caused by *P. capsici* expressing plasma membrane marker fused with mCherry. The arrow marks an aperture in the cell wall that is not accompanied by haustoria presence, h-haustorium, scale bar 2 μm, cell wall stained with aniline blue. Chloroplasts labeled in green.

**Supplemental Figure S4.**
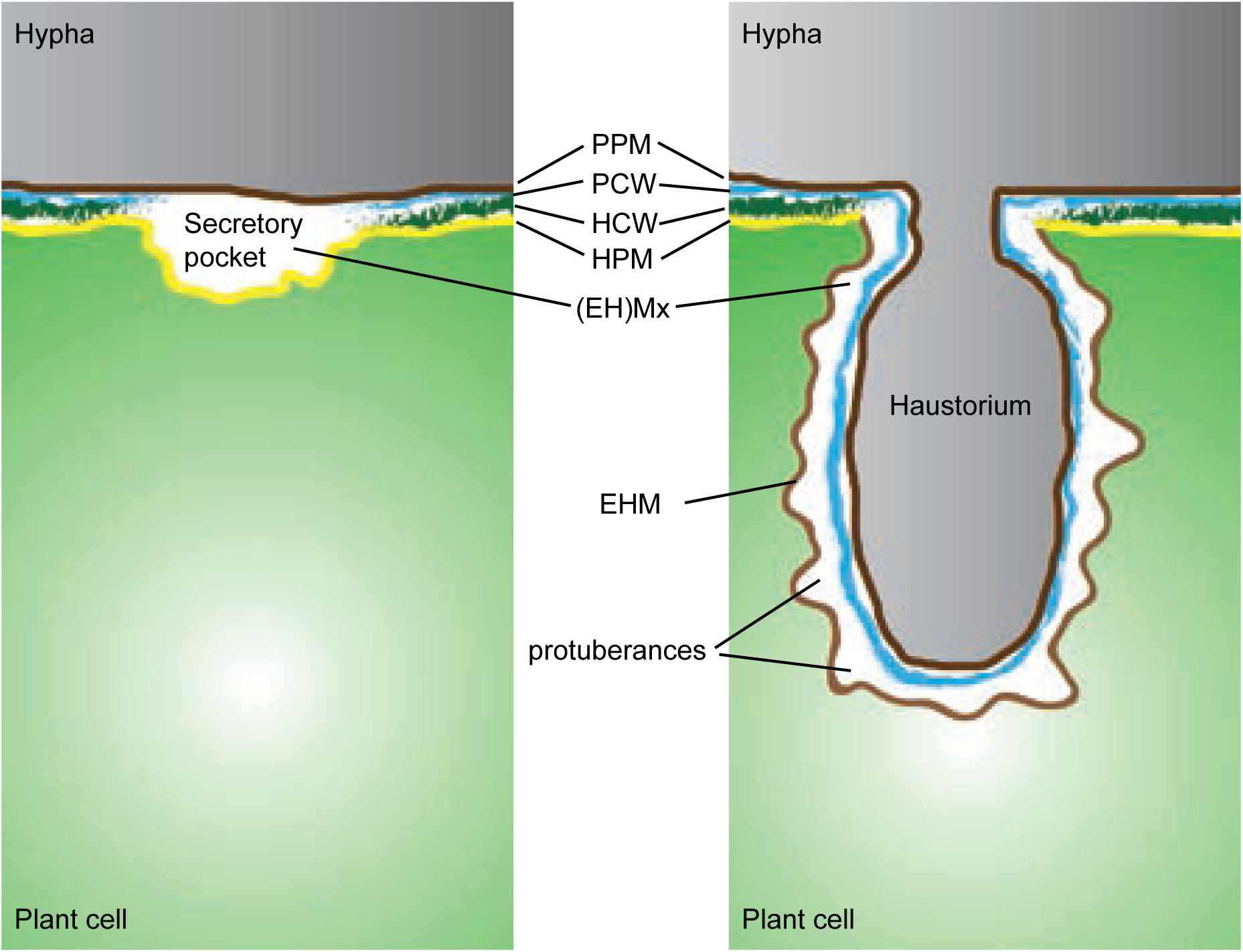
Schematic representation of the plant–pathogen interface: a secretory pocket which is the matrix filled with Phytophthora secretory material surrounded by the invaginated plant plasma membrane vs organization of a mature haustorium (based on the micrograph from Enkerli *et al*., 1997). Phytophthora hypha in gray, penetrated plant cell in green, HCW-Host cell wall, HPM-Host plasma membrane, PCW-Phytophthora cell wall, PPM-Phytophthora plasma membrane, (EH)Mx-(Extrahaustorial) matrix with its formed protuberances, EHM-Extrahaustorial membrane is convoluted.

**Supplemental Figure S5.**
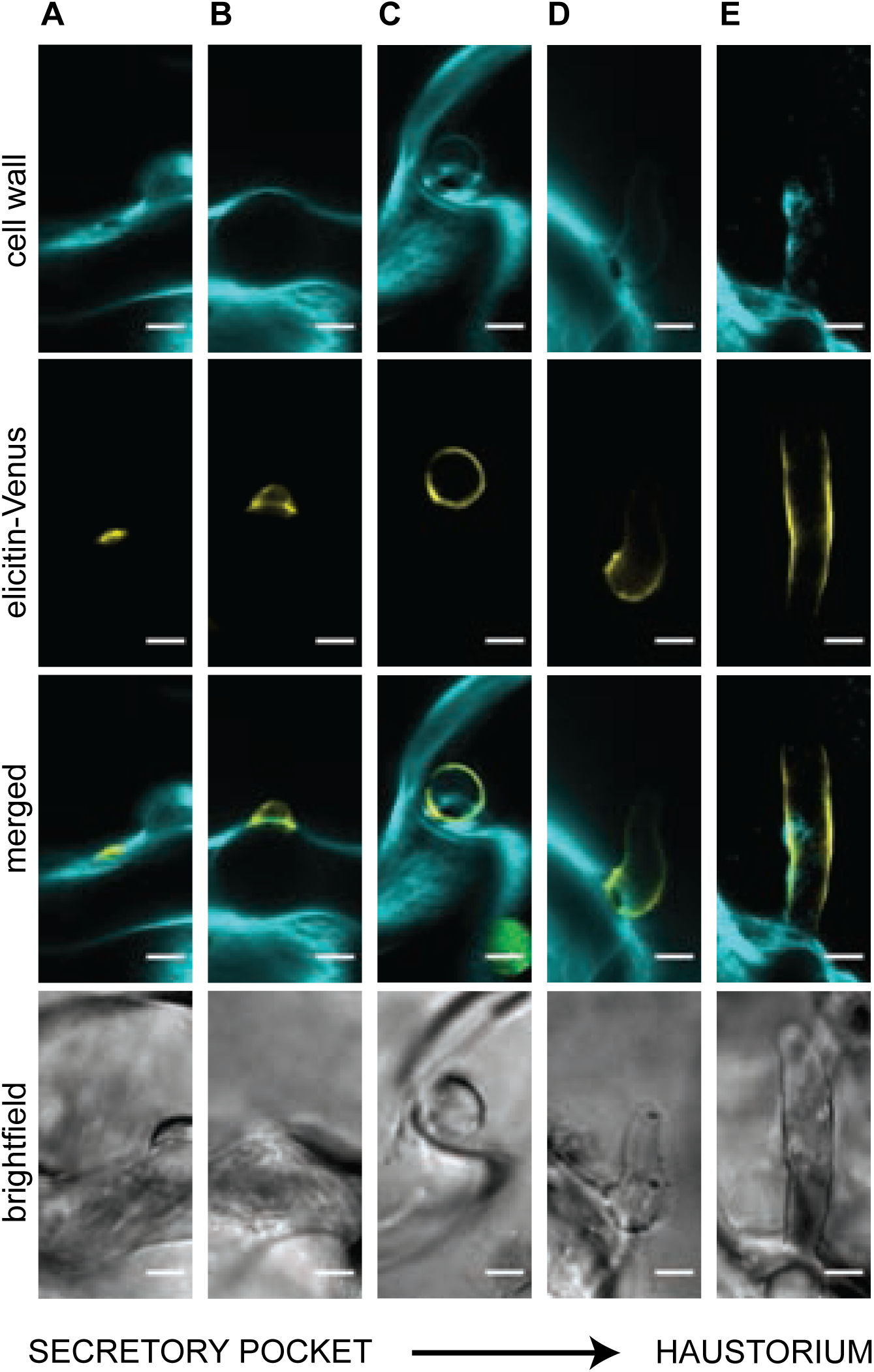
Development of sites for *P. capsici* protein secretion during *N. benthami ana* infection. Confocal microscopy pictures of elicitin-Venus expressed and secreted by *P. capsici* to (**A**) the secretory pocket, (**B**) and (**C**) intermediate growth stages between the secretory pocket and the haustorium, (**D**) mature haustorium. (**E**) Confocal microscopy pictures (maximal projection z-stack of 6 slices) of haustorium that expanded to the length of 17μm. Chloroplasts labeled in green. Cell wall stained with calcofluor white, scale bars 3μm.

**Supplemental Figure S6.**
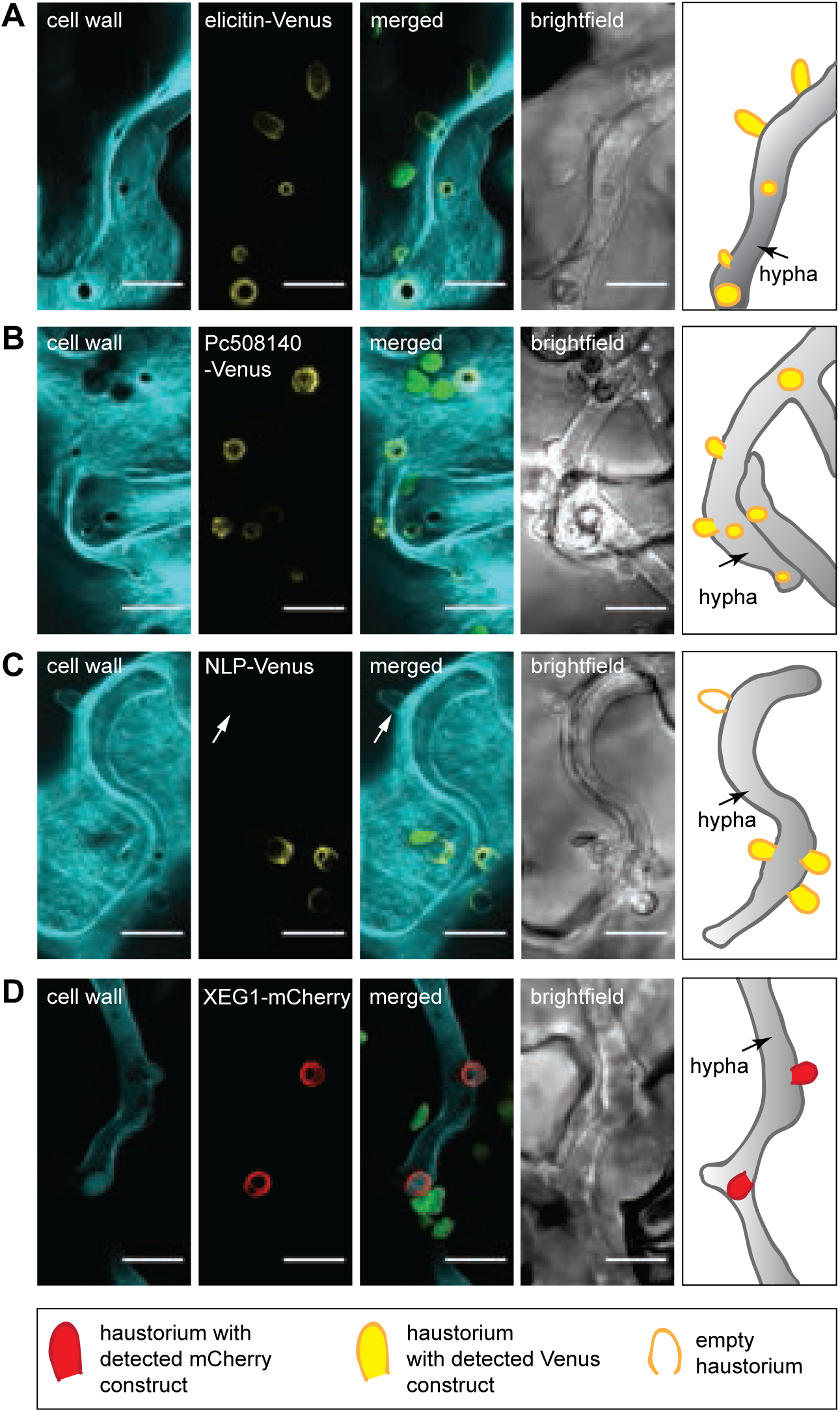
Apoplastic *P. capsici* proteins are delivered to haustoria during infection. Representa tive confocal microscopy pictures (maximal projection z-stack) of proteins expressed by *P. capsici* and accumulated at its haustoria during infection: (**A**) elicitin-Venus (12 slices), (**B**) Pc508140-Venus (15 slices), (**C**) NLP-Venus (12 slices), empty haustorium marked with an arrow, (**D**) XEG1-mCherry (11 slices). Cell wall stained with calcofluor white (**A**-**C**) or aniline blue (**D**). Chloroplasts labeled in green, scale bars 10 μm. All pictures accompanied by the schematic diagrams on the right side.

**Supplemental Figure S7.**
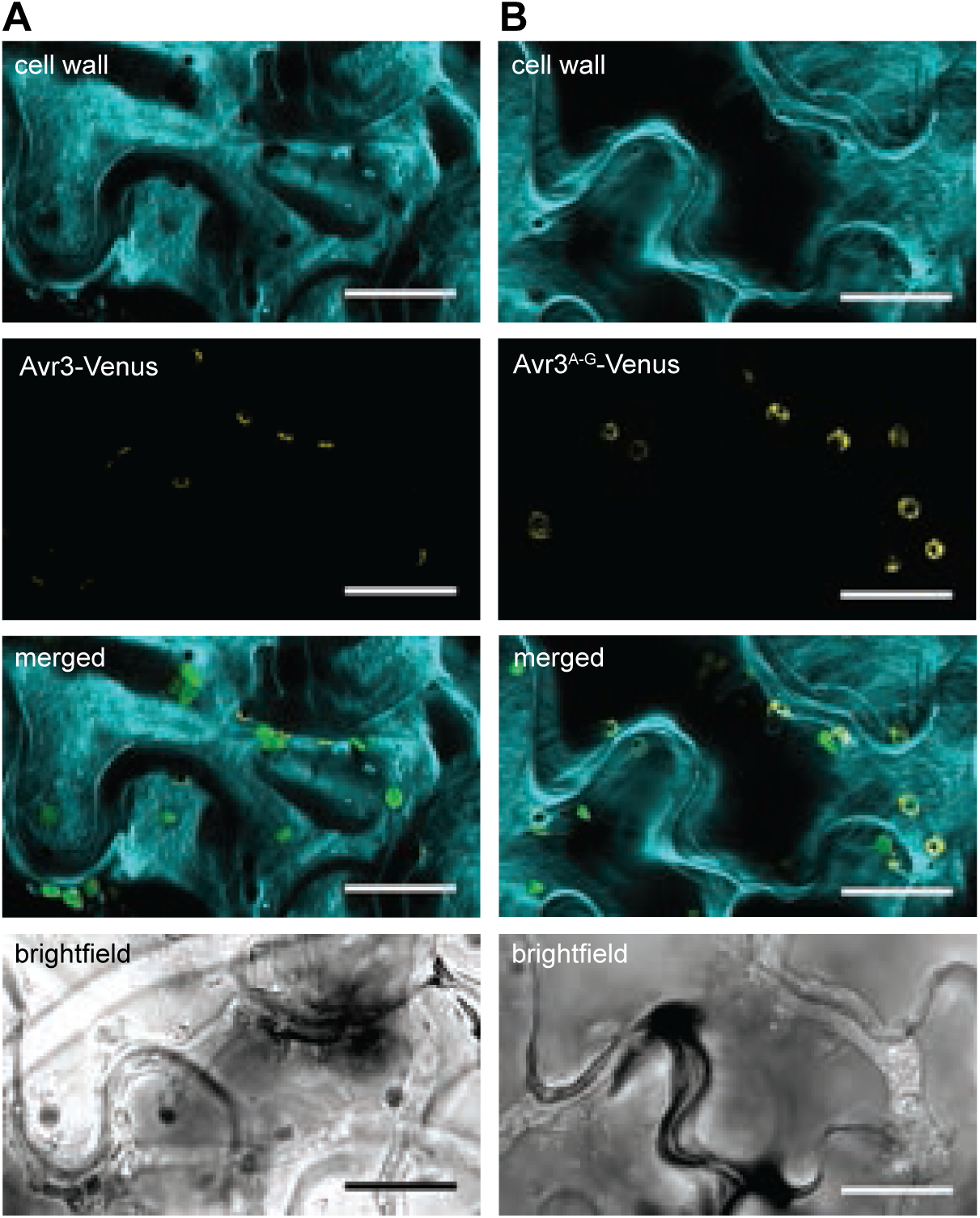
Avr3-Venus and its RXLR-EER motif mutant show different accumulation patterns at haustoria. Confocal microscopy pictures of *P. capsici* hyphae expressing (**A**) Avr3-Venus, the picture is a maximal projection z-stack of 7 slices, (**B**) Avr3-Venus with RxLR-EER fragment mutated to alanine-glycine residues. The picture is a maximal projection z-stack of 18 slices. Chloroplasts labeled in green. Scale bars 20 μm.

**Supplemental Figure S8.**
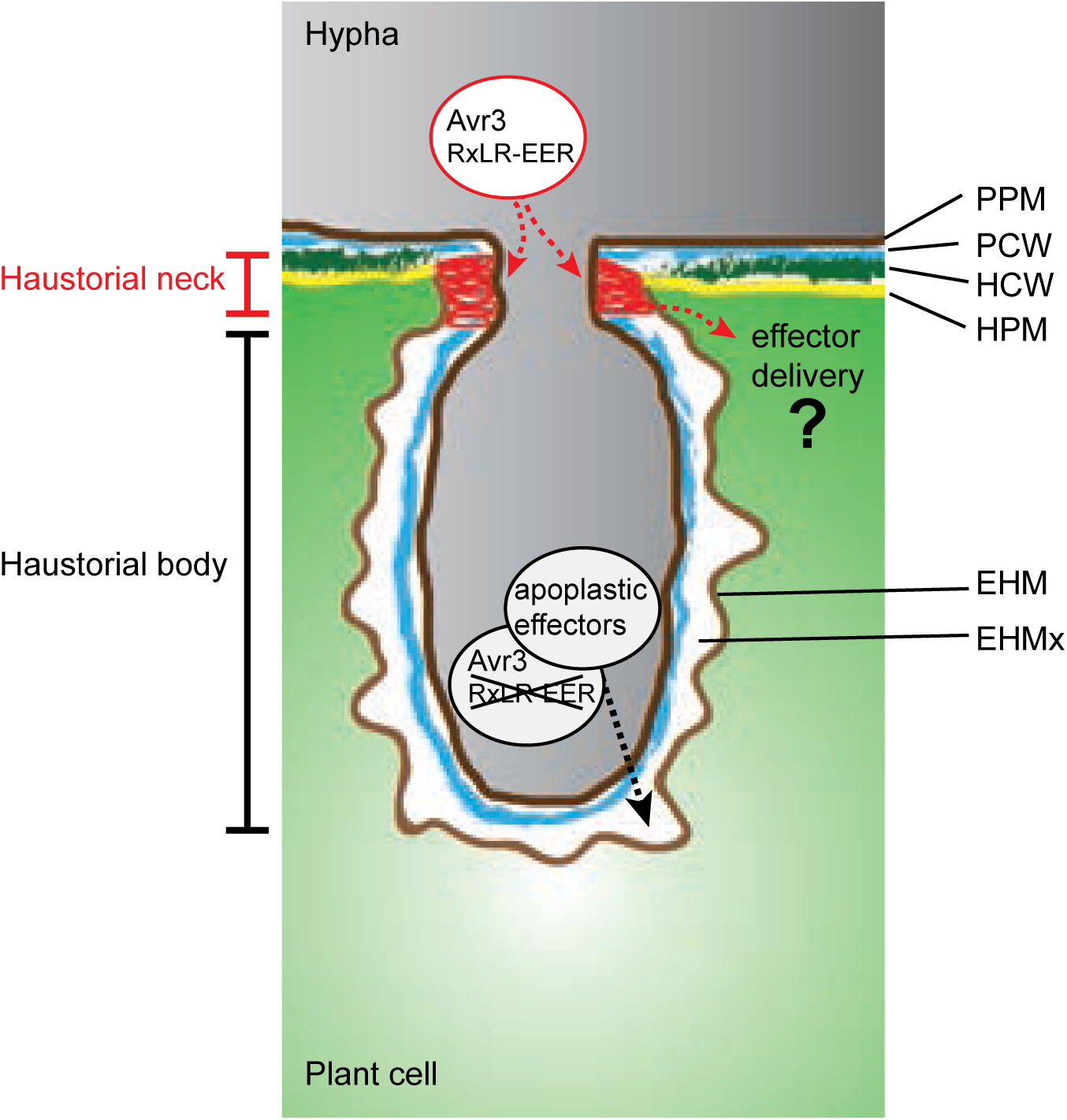
Model of effector secretion by *Phytophthora capsici*. The illustration shows two distinct regions where effectors accumulate at the haustorial interface. The RxLR effector Avr3 is localized at the haustorial neck region, suggesting its involvement in Avr3 translocation into the host cell. In contrast, a mutation in the RxLR-EER fragment of Avr3 redirects it to the EHMx, where apoplastic effectors are also secreted. The Phytophthora hyphae are shown in gray, the infected plant cell is depicted in green. HCW-Host cell wall, HPM-Host plasma membrane, PCW-Phytophthora cell wall, PPM-Phytophthora plasma membrane, EHMx-Extrahaustorial matrix, EHM-Extrahaustorial membrane.

**Supplemental Figure S9.**
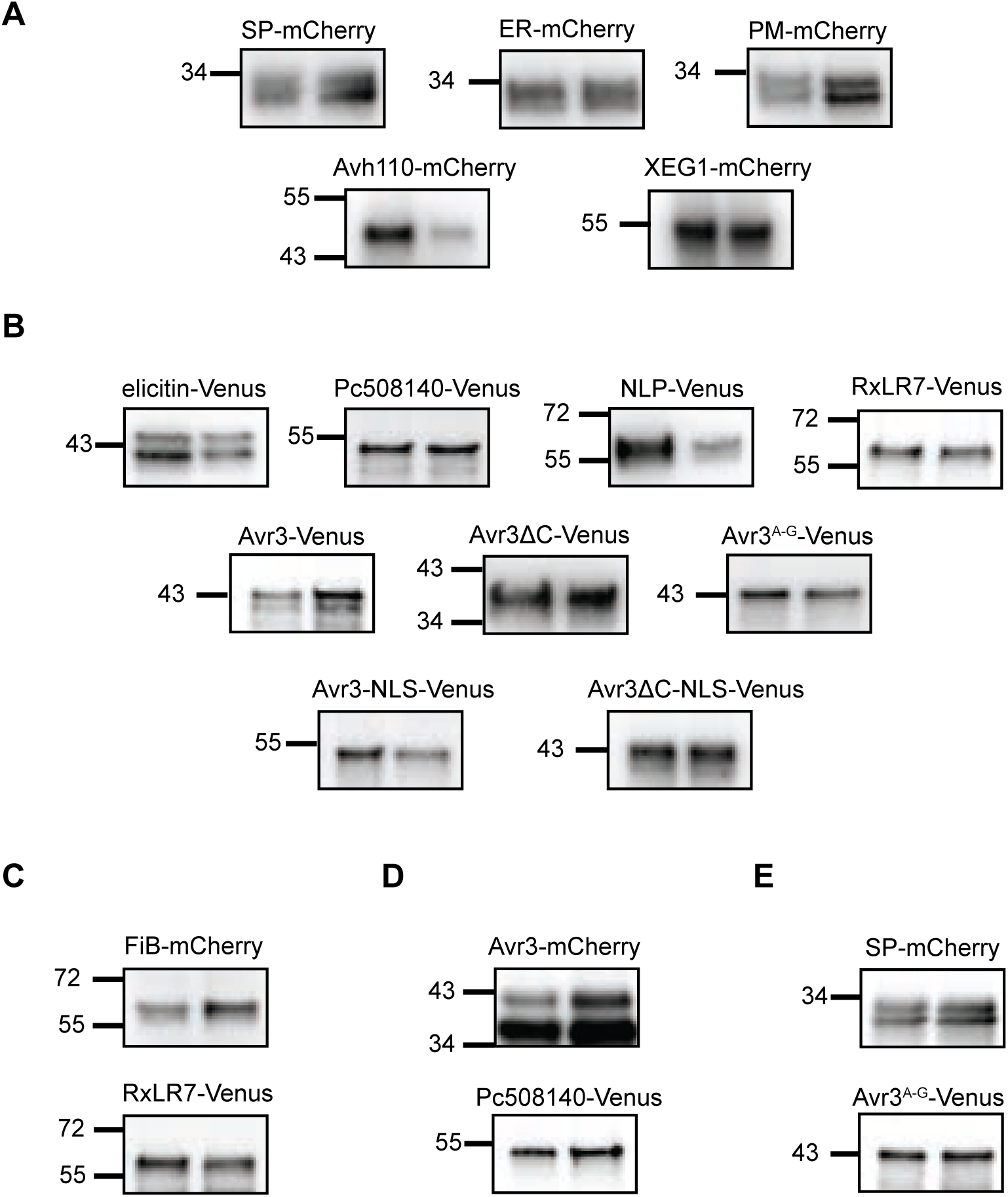
The expression of mCherry and Venus fusion proteins by *P. capsici* transformants was verified through immunoprecipitation from infected *N. benthamiana* tissue, followed by immunoblot analysis. For each construct, detection was carried out on samples infected by two indepen dent transient *P. capsici* transformants, with use of primary antibodies against (**A**) mCherry and (**B**) Venus. (**C**), (**D**), (**E**) Both antibody were used for detection of fusion mCherry and Venus proteins in double *P. capsici* transformants. Protein size markers are shown in kilodaltons. The expected protein sizes as follows: SP-mCherry 29 kDa, ER-mCherry 29 kDa, PM-mCherry 28kDa, Avh110-mCherry 48kDa, XEG1-mCherry 52 kDa, elicitin-Venus-FLAG 42kDa, Pc508140-Venus-FLAG 49 kDa, NLP-Venus-FLAG 57 kDa, RxLR7-Venus-FLAG 57 kDa, Avr3-Venus-FLAG 44 kDa, Avr3ΔC-Venus-FLAG 37kDa, Avr3A-G-Venus 44 kDa, Avr3-NLS-Venus-FLAG 49 kDa, Avr3ΔC-NLS-Venus-FLAG 42 kDa, FiB-mCherry 54 kDa, Avr3-mCherry 40 kDa.

